# Structural Basis of Redox-Coupled CO_2_ hydration by the Cyanobacterial NDH-1MS’ complex

**DOI:** 10.64898/2026.08.03.742399

**Authors:** Yat Kei Lo, Florian Ruoß, Jacqueline Thiemann, Marc Nowaczyk, Jan M. Schuller

**Affiliations:** Philipps-University Marburg, Department of Chemistry and SYNMIKRO Research Center, Marburg, Germany; Department of Chemistry for Life Sciences, Uppsala University, Uppsala, Sweden; Department of Biochemistry, University of Rostock, Albert-Einstein.Str. 3, 18059 Rostock, Germany; Plant Biochemistry, Faculty of Biology and Biotechnology, Ruhr-University Bochum, Bochum, Germany; Department Life, Light, Matter, University of Rostock, 18059 Rostock, Germany; Microbes-for-Climate (M4C) Cluster of Excellence, Marburg, Germany

## Abstract

The NAD(P)H dehydrogenase-like complex, NDH-1MS’, is a constitutively expressed CO_2_-concentrating mechanism critical for sustaining CO_2_ fixation in cyanobacteria. This complex has been proposed to function as a vectorial carbonic anhydrase and accumulates intracellular bicarbonate against chemical equilibrium by coupling CO_2_ hydration to the photosynthetic cyclic electron flow. Using cryo-electron microscopy, we determined the structure of NDH-1MS’ from *Thermosynechococcus vestitus*. Contrary to its inducible counterpart NDH-1MS, our data suggest that a candidate CO_2_ hydration site may reside in the carbon-concentrating subunit CupB instead of the CupB-NdhF4 interface previously proposed. The active-site region partially resembles metal ion-independent iota-carbonic anhydrases and may bind CO_2_ or bicarbonate. We further observe features consistent with a Grotthuss-type proton transfer network connecting the active-site region to the antiporter-like subunits NdhF4 and NdhD4; however, unlike complex I, the proton export channel in NdhF4 appears to be blocked by bulky hydrophobic amino acids. Structural comparison between the oxygenic photosynthesis-specific subunit NdhV-bound state and dissociated state further reveals a correlation between plastoquinone stability and NdhV association. Taken together, we propose a revised working model in which NDH-1MS’ may function as a metal ion-independent vectorial carbonic anhydrase, with CO_2_ hydration coupled to proton transfer events in the antiporter-like subunits across the thylakoid membrane.

**Significance:** Photosynthetic complex I-like CO_2_-uptake systems are central to cyanobacterial carbon fixation, yet the molecular basis for coupling CO_2_ hydration to electron transfer has remained unresolved. Here, structures of NDH-1MS’ in NdhV-bound and NdhV-free states revise the architectural framework for catalysis and coupling in the constitutive CO_2_-uptake module. The data identify a CupB-centered candidate active-site region, reveal a non-canonical terminal antiporter-like subunit with an apparently blocked P-side exit, and link NdhV association to altered plastoquinone stability. Together, these findings refine current models for vectorial carbonic anhydrase activity in cyanobacterial NDH-1 and provide a structural basis for understanding how photosynthetic redox chemistry may be coupled to inorganic carbon concentration.

## Introduction

Cyanobacteria, as evolutionary precursors of chloroplasts, alongside algae and plants, constitute the only organisms capable of oxygenic photosynthesis, a process fundamental to sustaining terrestrial life^1^. The Calvin–Benson cycle fixes atmospheric CO_2_ through carboxylation of ribulose-1,5-bisphosphate (RuBP) by the key enzyme ribulose-1,5-bisphosphate carboxylase-oxygenase (RuBisCO). However, the enzyme’s dual activity also leads to RuBP oxygenation, initiating photorespiration^2,3^, which detrimentally reduces photosynthetic efficiency by 20–50%^4^. To mitigate this limitation and adapt to environments with low environmental CO_2_, cyanobacteria have evolved sophisticated CO_2_-concentrating mechanisms (CCMs) that elevate intracellular bicarbonate levels and concentrate CO_2_ near RuBisCO, optimizing carboxylation and suppressing photorespiration^5,6^.

A major challenge faced by aquatic photoautotrophs like cyanobacteria lies in the limited bioavailability of dissolved inorganic carbon (DIC), governed by slow diffusion in aqueous environments and the pH-dependent equilibrium between CO_2_ and bicarbonate (HCO₃⁻)^7^. These two forms of inorganic carbon differ markedly in membrane permeability: uncharged CO_2_ readily diffuses across the cell membrane, whereas the charged bicarbonate ion requires specific transporter proteins for cellular uptake. Consequently, cyanobacteria have evolved diverse uptake systems tailored to environmental pH and inorganic carbon speciation. Under neutral to alkaline conditions, active transport systems facilitate bicarbonate uptake via three characterized transporters: the ATP-binding cassette (ABC)-type transporter BCT1^8^, and two Na⁺-dependent symporters, SbtA (high affinity)^9^ and BicA (low affinity)^9^. Conversely, in acidic conditions where dissolved CO_2_ predominates, specialized CO_2_ uptake systems convert CO_2_ into bicarbonate intracellularly^10–12^, thereby trapping it in an ionized, less membrane-permeable form.

At the core of this CO_2_ uptake strategy lies the functional versatility and modularity of photosynthetic NADH dehydrogenase-like complex I (NDH-1)^13^. The canonical NDH-1L complex, structurally homologous to respiratory complex I, operates as a ferredoxin:plastoquinone (Fd:PQ) oxidoreductase and plays a pivotal role in cyclic electron flow (CEF) around photosystem I^14^. It accepts electrons from reduced ferredoxin and transduces this redox potential into proton motive force via proton transport across the thylakoid membrane. This reaction enhances the ATP/NADPH balance under fluctuating light conditions^15^. A defining feature of NDH-1L is its incorporation of oxygenic photosynthesis-specific (OPS) subunits, such as NdhS and NdhV, that replace the NADH dehydrogenase module NuoEFG and instead facilitate ferredoxin binding^16^.

Specialized isoforms such as NDH-1MS and NDH-1MS’ retain the conserved electron-translocating membrane arm of NDH-1L but are functionally reconfigured through the modular exchange of terminal subunits, repurposing redox energy for vectorial hydration of CO_2_^17,18^. In these complexes the canonical antiporter-like subunits (ALS) NdhD1 and NdhF1 are replaced by paralogous pairs: NdhD3/NdhF3 in NDH-1MS and NdhD4/NdhF4 in NDH-1MS’. These isoforms stably associate with carbonic anhydrase-like subunits CupA/CupS or CupB, forming energy-coupled CO_2_ hydration modules. Expression studies and isotopic CO_2_ labeling experiments indicate that NDH-1MS functions as a high-affinity inducible system under low-CO_2_ conditions, while NDH-1MS’ constitutes a low-affinity, constitutively expressed form whose function is to capture CO_2_ leaked out of carboxysomes^10,19^. Notably, although CupA and CupB lack canonical Zn²⁺-binding motifs typical of carbonic anhydrases, genetic and physiological data as well as membrane-inlet mass spectrometry of mutants support their function in unidirectional CO_2_ hydration to HCO₃⁻^19–21^. This reaction is thought to be rendered vectorial by coupling to the proton-translocating activity of NDH-1, thereby preventing cytosolic equilibration and loss of CO_2_ via passive diffusion^17^.

Cryo-EM analysis of the NDH-1MS complex from *T. vestitus* revealed that CupA and CupS are positioned at the distal end of the membrane arm, forming a distinctive U-shaped architecture^17,22^. The catalytic site was proposed to reside at the CupA–NdhF3 interface and coordinates a Zn²⁺ ion via conserved residues (CupA His130, Arg135; NdhF3 Arg37, Glu114, Tyr41), consistent with a functional CA-like site. Although NdhF3 shares structural homology with Mrp-type antiporters, it lacks the conserved charged residues required for transmembrane proton translocation, and molecular dynamics simulations suggest it is incapable of supporting a functional proton-conducting tunnel^17^. It is proposed to form a putative intramembrane gas conduit connecting the bilayer interior to the CupA active site, offering a mechanistic basis for CO_2_ channeling to the catalytic center. In contrast, NdhD3 retains key antiporter-like features and may mediate proton translocation through a yet-unresolved structured network of water molecules or lipid headgroups, potentially forming transient water wires to link NdhD3 to the CupA active site, but such a pathway is not resolved in current cryo-EM reconstructions^17^. Thus, how proton translocation via NdhD3 is structurally and functionally coupled to CupA to enable the redox-driven vectorial hydration remains an open and critical question.

Here, we solved the structures of NDH-1MS’ in the NdhV-bound state and in the NdhV-dissociated state using cryogenic-electron microscopy. Our high-resolution models reveal an unexpected alternative active-site candidate within CupB and features consistent with a Grotthuss-type proton transfer network linking this region to the ALS. Furthermore, structural comparisons identified key differences at the plastoquinone pocket between the two states, and unique features in NDH-1MS’ ALS. Integrating the findings, we propose a revised working model for how CO_2_ hydration may be coupled to proton transfer events in the terminal antiporter-like region of NDH-1MS’.

## Results

### Overall structure and cofactors

We purified the NDH-1MS’ complex from *T. vestitus* BP-1 using a twin-strep tag on the CupB subunit and determined its cryo-EM structure in GDN at 2.3 Å resolution (Fig. 1a, Supplementary Fig. 1, 2, Supplementary Table 1). The map resolved all core subunits (NdhA– C, NdhE, NdhG–K, NdhM/N) and the oxygenic photosynthesis-specific (OPS) subunits (NdhL/O/S), along with the carbon-concentrating NDH-1S’ module (NdhD4/F4, CupB). Local resolution decreases toward the NDH-1S’ module, indicating higher flexibility in this region compared to the core. To improve map quality, we performed local refinement of NdhF4 and CupB, yielding a focused reconstruction at 2.4 Å (Supplementary Fig. 2, Supplementary Table 1). Previous NDH-1 structures were determined either without NdhV and ferredoxin (Fd) or with both components added in excess^14,17,23–25^. In contrast, focused classification of the NdhV binding site enabled us to isolate a subpopulation (32%) of NDH-1MS’ containing NdhV but lacking Fd, which we resolved at 2.7 Å (Fig. 1a, Supplementary Fig. 2, Supplementary Table 1). We hereafter refer to this subpopulation as NDH-1MS’–NdhV. This finding demonstrates, for the present preparation, that NdhV can associate transiently and independently of Fd. Unless otherwise specified, our structural analyses are based on the higher-resolution NDH-1MS’ map.

**Figure 1.**
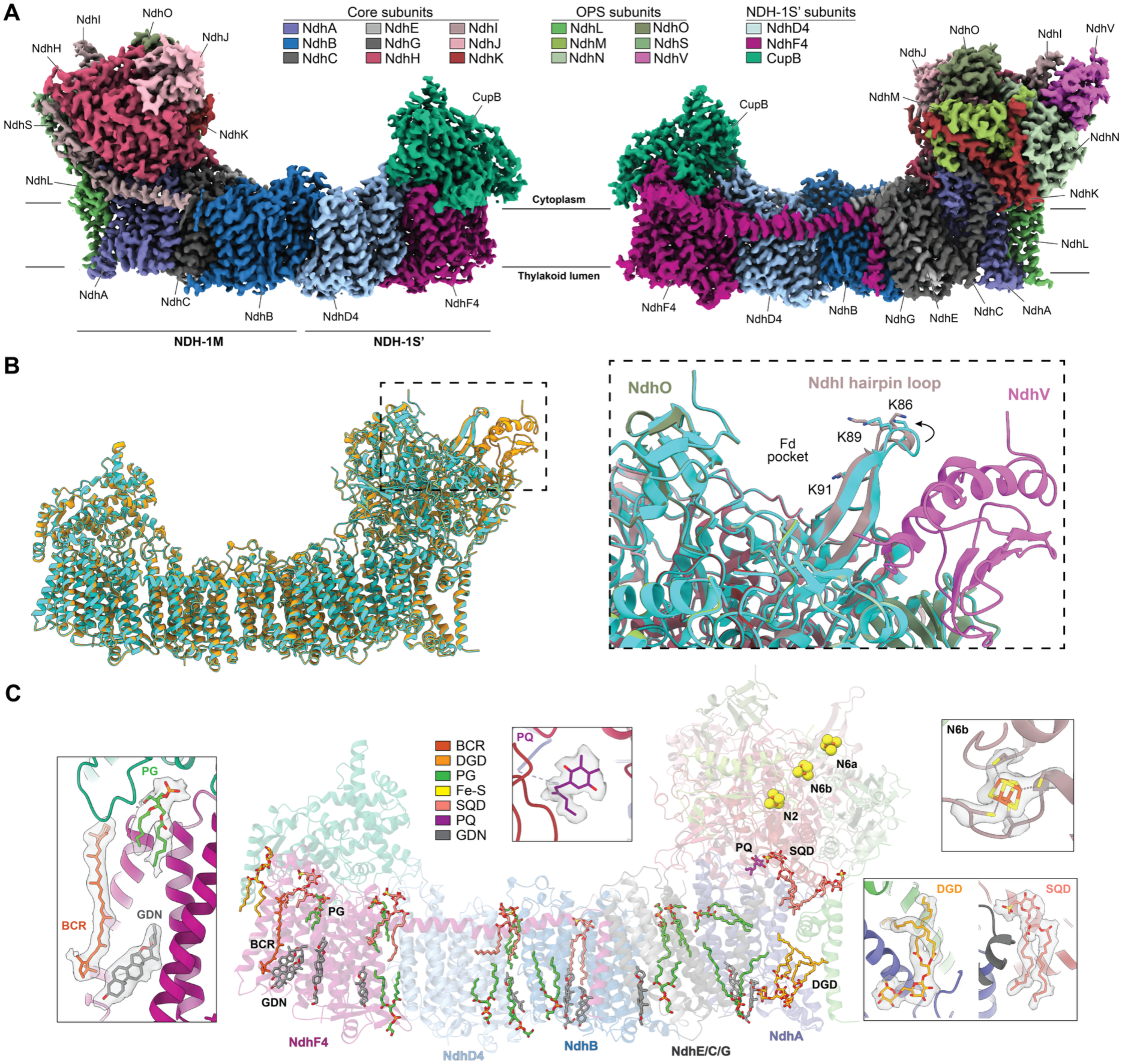
Structural overview and cofactors. **A** Front view of NDH-1MS’ consensus volume (left) and back view of NDH-1MS’-NdhV volume (right). **B** Left: Superposition of NDH-1MS’ structural model (cyan) on NDH-1MS’-NdhV (orange). Right: Zoom-in view of the Fd binding pocket. NDH-1MS’-NdhV is colored as in panel A; NDH-1MS’ in cyan. **C** Overview of cofactors and ligands modeled on NDH-1MS’ consensus map. Representative density is shown in inserts. BCR: β-carotene; SQD: sulfoquinovosyl diacylglycerol; DGD: digalactosyl diacylglycerol; PQ: plastoquinone; PG: phosphatidylglycerol; GDN: glycol-diosgenin; Fe-S: 4Fe-4S cluster.

NDH-1MS’ closely resembles NDH-1MS^17^, with CupB positioned directly above NdhF4 in nearly the same orientation as CupA in NDH-1MS (Fig. 1a, 2a). NDH-1MS’–NdhV also matches the structure of NDH-1L in the NdhV–Fd co-bound state^25^, where NdhV binds to the apex of the peripheral arm and interacts with NdhI/N/S. No major structural difference was observed between NDH-1MS’ and NDH-1MS’–NdhV (RMSD = 0.26 Å), except for a slight displacement of the NdhI hairpin loop towards the Fd-binding pocket in the NdhV-bound state (Fig. 1b). Similar conformational changes stabilize Fd binding in NDH-1L when both NdhV and Fd are present^23^. Although it remains unclear whether this shift is driven by Fd or NdhV, our structure suggests that NdhV alone can induce hairpin loop repositioning, thereby priming the site for Fd association.

**Figure 2.**
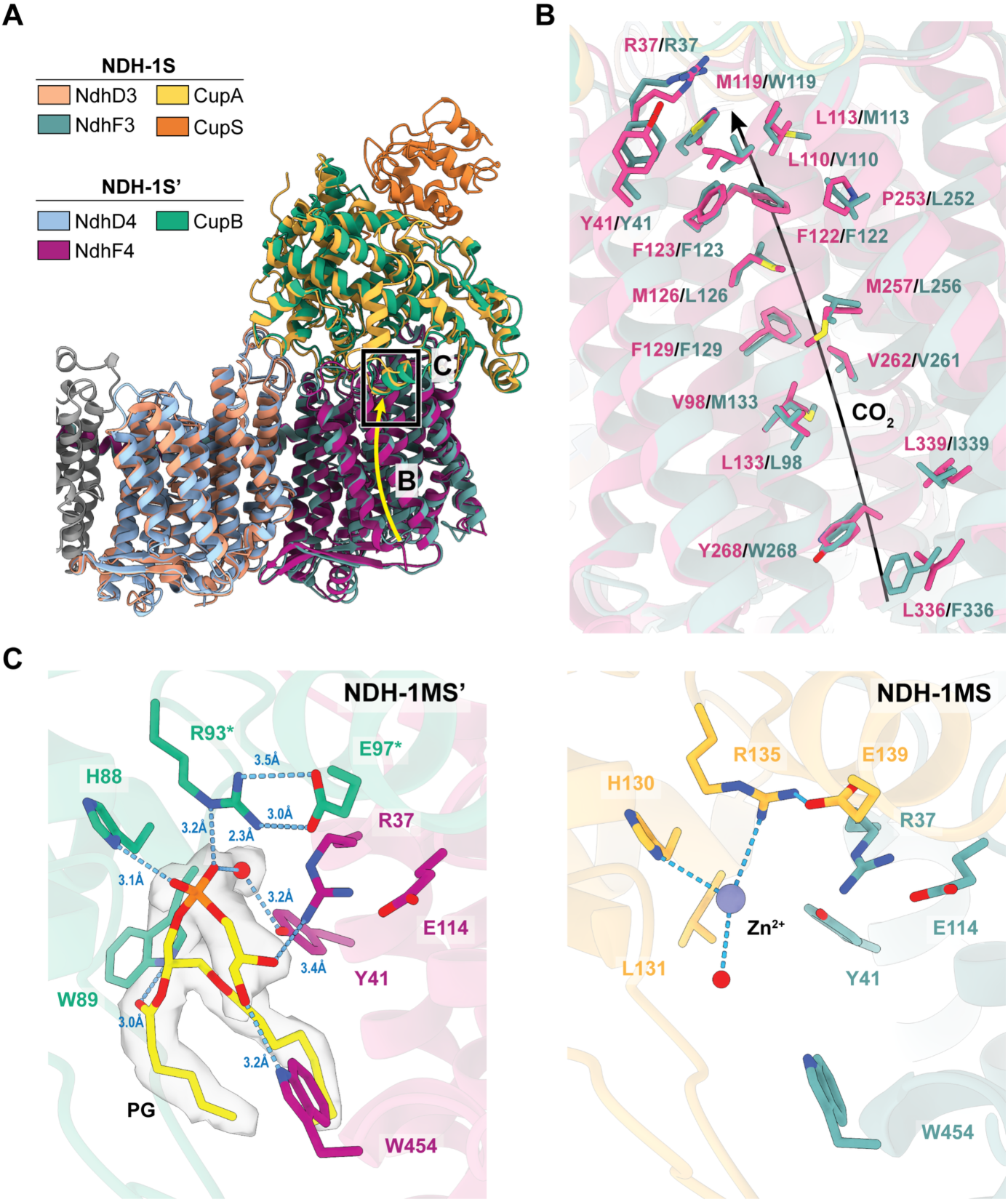
Carbon-concentrating module of NDH-1S/S’. **A** Superposition of NDH-1S (PDB 6TJV) on NDH-1S’. Yellow arrows and black square indicate the CO_2_ tunnel and position of active site previously proposed in NDH-1S. **B** Comparison of key residues lining the putative CO_2_ tunnel (depicted by arrow) between NDH-1S and NDH-1S’. **C** Side-by-side comparison of the proposed Zinc-coordinating active site between the two structures. PG Density of NDH-1MS’ focused refinement are displayed at 4 σ. Blue dashes depict polar interactions. Asterisk (*) indicates residues strictly conserved among 150 homologue as calculated by the Consurf server^64^.

Our cryo-EM density also resolved several thylakoid membrane lipids and essential cofactors (Fig. 1c), including all three 4Fe–4S clusters along the peripheral arm and a plastoquinone (PQ) molecule at the ‘medium’ binding site^26^ just above the membrane plane. Lipids line the membrane arm and cluster at subunit interfaces between NdhA/C/K/L, NdhB/D4, and NdhD4/F4, resembling those observed in NDH-1L and NDH-1MS. These clusters likely stabilize the membrane arm, as proposed for other NDH-1 complexes^23,25^. In contrast to NDH-1L^23^, no carotenoid was found between NdhD4 and NdhF4, suggesting that carotenoid binding at this site is specific to NDH-1L and depends on NdhP/Q, which are absent in NDH-1MS and NDH-1MS’. Instead, NdhF4 harbors a single carotenoid at its peripheral end, directly exposed to the detergent micelle (Fig. 1c). This carotenoid may stabilize the transmembrane helices of NdhF4 or reinforce its interaction with CupB, functioning as a molecular bridge as in NDH-1MS. However, unlike NDH-1MS^17^, NdhF4 lacked any structurally resolvable chlorophyll.

### Active site and assembly of the carbon-concentrating module

The architecture of NDH-1S’ closely resembles that of the NDH-1S module. Previous structural analyses and molecular dynamics (MD) simulations of NDH-1MS identified a potential CO_2_ tunnel in NdhF3 that connects the thylakoid lumen to the proposed active site at the CupA–NdhF3 interface^17^. Superimposition of NDH-1S on NDH-1S’ revealed that the non-polar residues forming this tunnel were conserved, suggesting that NdhF4 may possess a similar potential gas tunnel (Fig. 2a, b). In NDH-1S, the proposed active site consists of a zinc ion coordinated by CupA residues His130 and Arg135, which form a salt bridge with Glu139. NdhF3 residues Arg31, Tyr41, and Glu114 further enclose the active site (Fig. 2c). CupB and NdhF4 also contain these conserved residues (CupB: Arg93, His88, Glu97; NdhF4: Arg31, Tyr41, Glu114). However, our high-resolution map revealed a well-defined phosphatidylglycerol (PG) density, including a fully resolved head group, occupying this position instead of a metal ion. The lipid engages in an extensive hydrogen-bonding network with CupB residues His88, Trp89, and Arg93, as well as NdhF4 residues Arg37 and Trp454 (Fig. 2c). Additionally, a water molecule coordinated by NdhF4 Tyr41 hydrogen bonds to the phosphate group of PG. Although we cannot rule out the possibility that NDH-1MS’ lost the zinc cofactor during purification, leaving an empty pocket that was later occupied by PG, this observation argues against the assignment of this position as the sole active site.

Our analysis also uncovered two previously unresolved densities (UNK1 and UNK2) located at the center of CupB (Fig. 3a). Using Caver^27^, we identified two tunnels: tunnel T1 connects UNK1 and UNK2 to the bulk solvent (Fig. 3b), while tunnel T2 extends toward the cytoplasmic half-channel of NdhF4 (Fig. 3c). Although the ligands could not be assigned with certainty, the geometry of UNK1 and UNK2 are compatible to a bicarbonate ion and a CO_2_ molecule, respectively (Fig. 3b). UNK1 is unlikely to represent a single ion or water molecule, as its density center lies more than 4 Å from the nearest probable coordinating residue. Aromatic residues predominantly enclose UNK1, with the putative bicarbonate ion coordinated by conserved residues Tyr47, Trp169, and Tyr240. Tyr47 additionally hydrogen bonds to a nearby water molecule adjacent to UNK1.

**Figure 3.**
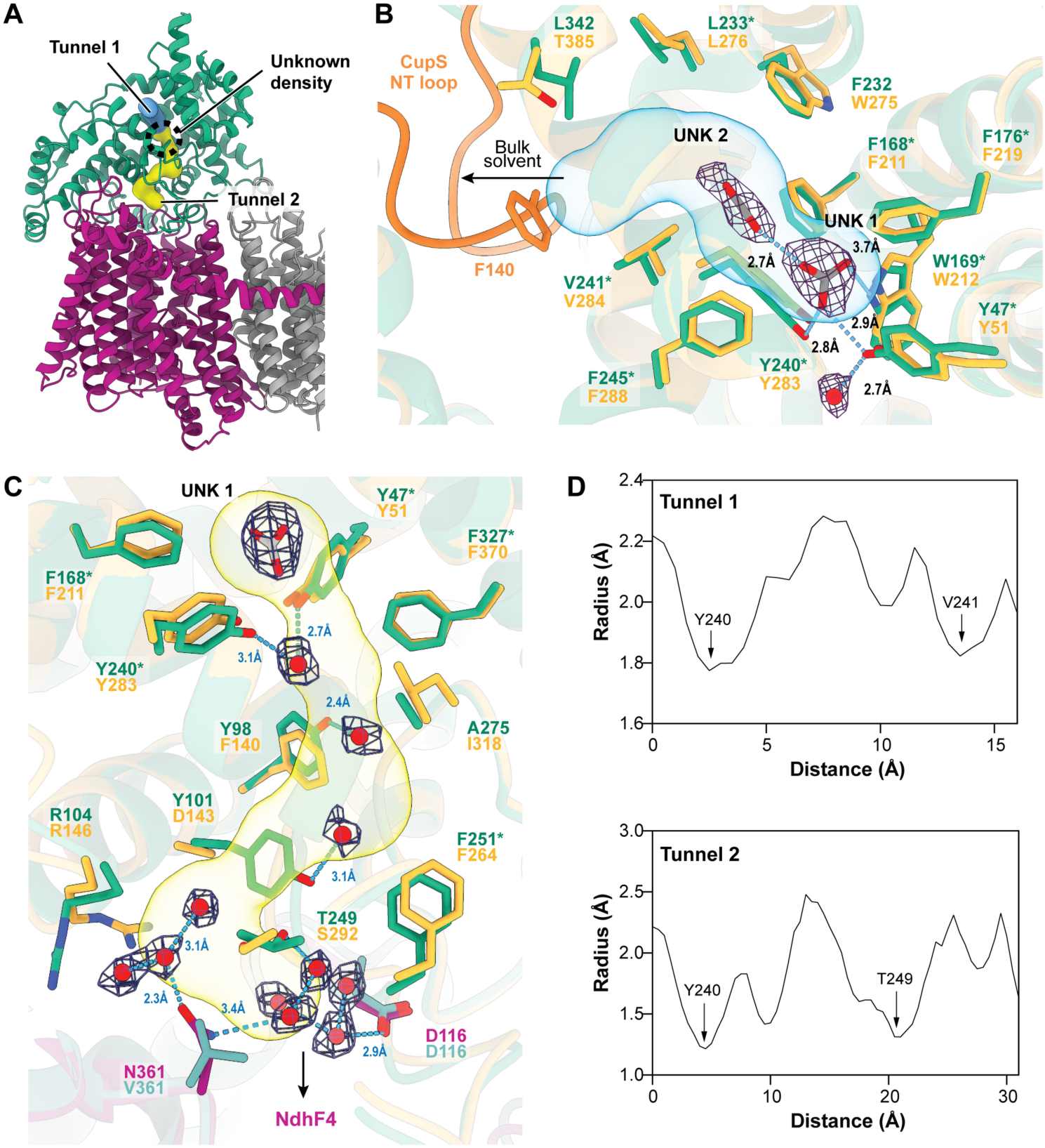
Unknown ligand densities in the NDH-1S’ module. **A** Position of the unknown densities and connecting tunnels predicted by Caver^27^ in NDH-1MS’. **B** Zoom-in view of the unknown densities and tunnel 1. CupB key residues around the unknown densities are shown and compared to CupA. Models are colored as in Fig. 2. Notice that the tunnel opening was blocked by CupS NT loop in NDH-1MS while remained open to the bulk solvent in NDH-1MS’. **C** Zoom-in view of tunnel 2 and residues lining the tunnel. The side chain of CupA residue D143 was truncated in the deposited model. **B, C** Ligand densities of the focused refinement are displayed at 6 σ. Blue dashes depict potential hydrogen-bonding network. Asterisk (*) indicates residues strictly conserved among 150 homologue as calculated by the Consurf server^64^. **D** Dimension of the predicted tunnels. Center of the UNK 1 density was defined as the tunnel origin.

Tunnel T1, extending toward the bulk solvent, is scaffolded by non-polar residues and has a predicted bottleneck radius of 1.8 Å (Fig. 3b, d). It is therefore predominantly hydrophobic and can accommodate CO_2_ (radius ∼1.6 Å). With conformational flexibility, the tunnel might also accommodate bicarbonate (radius ∼2.6 Å). Although most of the non-polar residues lining this tunnel are conserved in CupA, the tunnel opening in CupA is blocked by the N-terminal loop of CupS, suggesting that CupS may regulate ligand access (Fig. 3b). Tunnel T2 is similarly non-polar but more restricted, with a bottleneck radius of 1.2 Å (Fig. 3d). It is partially hydrated by structurally resolved waters hydrogen-bonded to CupB residues Tyr98, Tyr101, Tyr240, and Thr249, as well as NdhF4 residues Asp116 and Asn361 (Fig. 3c). While the precise function of this tunnel remains unclear, its narrow dimensions imply that it can accommodate water molecules but is unlikely to transport larger ligands such as bicarbonate without significant rearrangements. The resolved waters may instead form part of a water wire linking UNK1 to the N-side half-channel of NdhF4.

Despite the strong structural homology between NDH-1S and NDH-1S’ subunits (Fig. 2a), they are not interchangeable and cannot assemble into chimeric complexes^19,20^. The assembly of NdhD4/F4 and NdhD3/F3 is driven by hydrophobic interactions between their transmembrane helices, aided by a few polar side-chain interactions (Supplementary Table 2). While some polar contacts are conserved between NdhD4/F4 and NdhD3/F3, superimposition of NdhD3 on NDH-1S’ revealed steric clashes between their interfacing helices and with the axial helix of NdhF4 (Supplementary Fig. 3). By contrast, assembly of the CupA–NdhF3 and CupB–NdhF4 modules rely mainly on polar interactions. Comparative analysis highlighted several CupB–NdhF4-specific interactions absent in CupA–NdhF3, while association of CupA with NdhF4 is sterically hindered by severe clashes across the interface (Supplementary Table 2, Supplementary Fig. 3). These observations indicate that, although NDH-1S and NDH-1S’ share a highly similar overall architecture, subunit specificity is dictated by both side-chain geometry and unique polar interactions.

### Architecture of the plastoquinone chamber

The peripheral arm of NDH-1MS’ and NDH-1MS’–NdhV resembles the open state of bacterial respiratory complex I^26,28^ and exhibited all defining features except for the NdhC TM1-2 loop. These features include the partially disordered NdhA TM5-6 loop, the π-bulged NdhG TM3, the extended NdhH β1-2 loop, and the raised NdhK σ1-2 loop (Fig. 4a, b, Supplementary Fig. 4). In our structure, a plastoquinone (PQ) molecule resided within the hydrated quinone chamber, above the membrane plane, in a position comparable to that reported in *E. coli* complex I^26^ (Fig. 4a, Supplementary Fig. 4). Only the head group and a short segment of the PQ tail were resolved in both of our density maps, as in the PQ-bound state of NDH-1L^23^. The relatively weak PQ density compared to surrounding amino acids suggests that PQ is highly flexible, and our structures likely capture a transient PQ-bound state. The PQ entrance from the thylakoid membrane is tightly restricted by hydrophobic residues of NdhA, with an opening radius of 1.1 Å as calculated by Caver (Supplementary Fig. 5). Access to the terminal 4Fe– 4S cluster N2 is blocked by the extended NdhH β1-2 loop (Fig. 4b), also seen in NDH-1L (Supplementary Fig. 4). Overall, association of the distal NDH-1S’ module does not alter the quinone chamber, and the NDH-1MS’ peripheral arm retained the same architecture as other NDH-1 complexes^14,17,23–25^.

**Figure 4.**
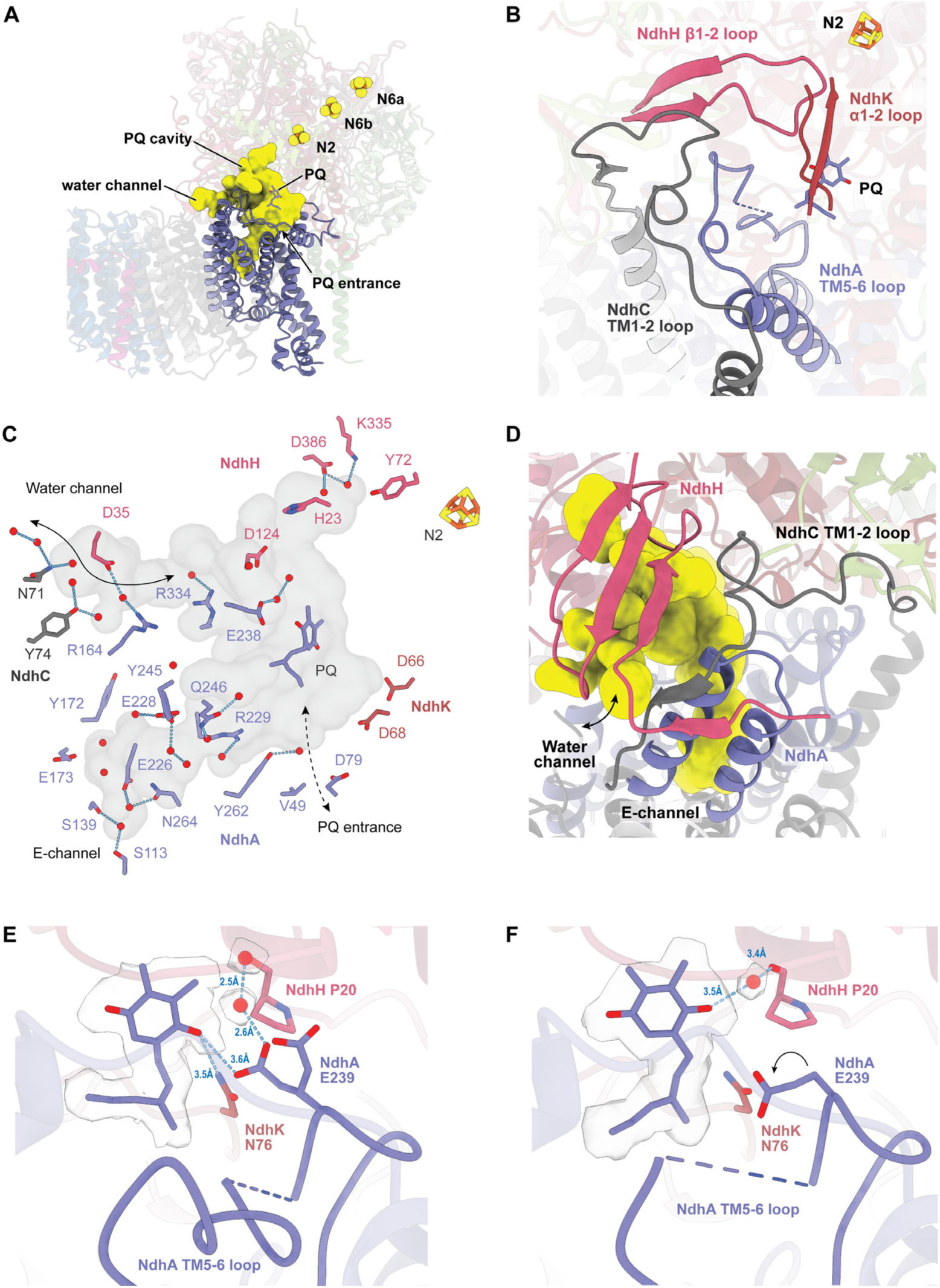
NDH-1MS’ quinone cavity. **A** Overview of the quinone cavity calculated by CASTp^65^ using a 1.4 Å radius probe. **B** Conformation of key loops enclosing the quinone chamber. **C** Architecture of the quinone cavity featuring conserved Glu and water network residues. **D** Opening of the proposed water channel for (de)hydration of the quinone cavity. Notice that the NdhC TM1-2 loop was fully stabilized by polar interactions between neighboring subunits (see Supplementary fig. 5). Coordination of PQ in **E)** NDH-1MS’ and **F)** NDH-1MS’-NdhV. Glu239 of NDH-1MS’ exhibited two side chain conformations. Ligand densities are displayed at 2.5 σ. Arrow indicated repositioning of Glu239 in the NdhV-bound state. Light blue dashes depict potential hydrogen-bonding network.

Contrary to the open state of complex I^26,29–31^, the NdhC TM1-2 loop (homologous to ND3/NuoA) was fully resolved in our structure (Fig. 4b, Supplementary Fig. 4). In *E. coli* complex I, the TM1-2 loop in the open state remains flexible to permit opening of the water channel gated by NuoCD (NdhH) that mediates hydration and dehydration of the quinone chamber^26^. Upon transition to the closed state, the TM1-2 loop becomes ordered, stabilized by polar interactions with NuoB/CD/H, while repositioning of NuoCD closes the water channel. A similar water channel is established in NDH-1MS’, where NdhH adopted the open conformation, while the NdhC TM1-2 loop is stabilized by polar interactions with NdhA/H/K (Fig. 4c, d, Supplementary Fig. 4, 5). Although NDH-1might alternate between open and closed states mirroring complex I given their homology and conserved coupling mechanisms, structures of NDH-1 resembling the closed state have not yet been determined. The unexpectedly stable NdhC TM1-2 loop, nonetheless, indicates that opening of the PQ chamber water channel might not require conformational changes of the TM1-2 loop in NDH-1.

A hallmark of the NdhV-bound state is destabilization of PQ. In the previously reported NDH-1L PQ-bound state^23^, the terminal backbone amide of Ala237 at the partially disordered TM5-6 loop of NdhA is proposed to hydrogen bond with PQ. However, this conformation is unfavorable in NDH-1MS’ since the TM5-6 loop would extend into the PQ pocket, potentially introducing steric clashes. Notably, based on NDH-1 structures available so far, the TM5-6 loop becomes stabilized only in the apo state irrespective of Fd or NdhV binding^14,17,24,25^. In NDH-1MS’, PQ hydrogen bonds to NdhK Asn76 and protonated Glu239 of NdhA, as predicted by PropKa^32^ (Fig. 4e, Supplementary Fig. 6). Glu239 also adopts an alternative conformation pointing away from PQ, indicating that the PQ–Glu239 interaction is likely to be transient. Upon NdhV binding, Glu239 rotates away, eliminating its interaction with PQ (Fig. 4f). This repositioning coincided with a slight shift of PQ toward the N2 cluster, where it was stabilized instead by a hydrogen bond from a water molecule. It remains inconclusive whether NdhV binding alone drives these conformational changes. Nonetheless, destabilization of PQ in the NdhV-bound state may facilitate PQ movement either into the deep binding site upon retraction of the NdhH loop or toward the shallow binding site.

### Proton translocation pathways

Provided the higher resolution in NDH-1MS’ and structural identity of the membrane arm between the two states, we focus our analysis on NDH-1MS’. The consensus volume revealed 303 structural water molecules along the membrane arm and an additional 30 in NdhF4 from the focused refinement (Fig. 5a, Supplementary Fig. 7). The overall hydration profile along the membrane arm resembles that of complex I^26,31,33–36^, except for NdhF4, which lacks connection to the thylakoid lumen (see below). From NdhD4 to NdhE, conserved central Lys, Glu, Asp, and His residues coordinate water molecules along the central membrane axis, establishing features consistent with a Grotthuss-type proton translocation network that potentially links NdhD4 TM12 Glu401 to NdhE TM3 Glu73. Several additional water molecules are stabilized by hydroxyl groups of polar residues and Tyr sidechains. The E-channel is similarly hydrated from the Glu cluster to NdhC Asp85. The connection between the central pathway and the E-channel is interrupted by a large hydrophobic gap between NdhE Glu73, NdhG Tyr66, and NdhC Asp85, consistent with the open state observed in complex I^30,31^. Although no continuous linkage from the E-channel to the thylakoid lumen was resolved, several water molecules appear near the putative channel exit.

**Figure 5.**
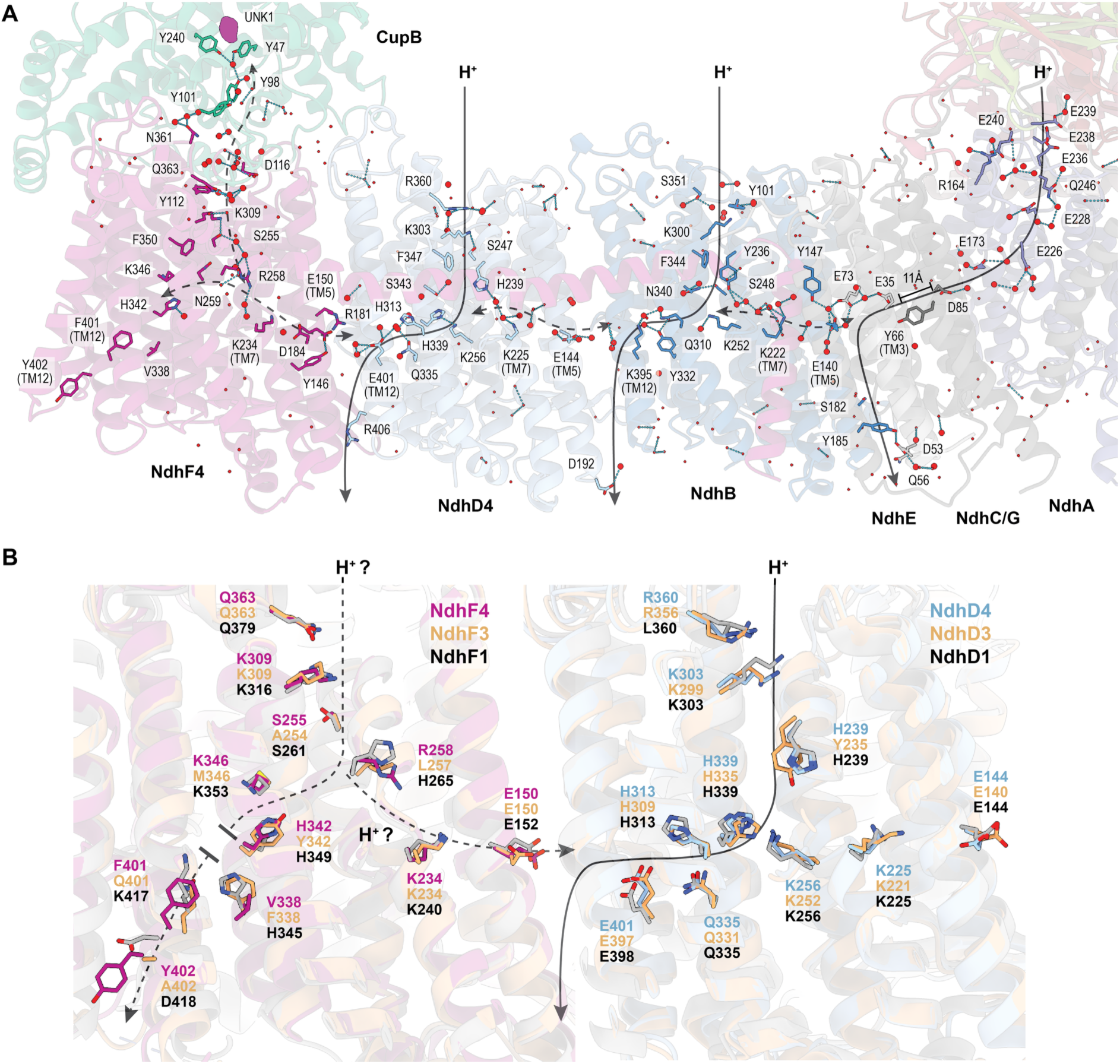
Water wire across the membrane arm. **A** Hydration profile of NDH-1MS’ membrane arm. Solid arrows indicate the proposed proton pumping pathway according to the “wave propagation” model. Dashed arrows indicate probable Grotthuss proton transfer network. Water within 5 Å of key proton network residues are depicted by red spheres. Water along the membrane arm but beyond 5 Å are depicted by red dots. Water molecules in NdhF4 and CupB were modelled on the focus refined volume while other water molecules were modelled on the consensus volume. The unknown density (UNK1) in CupA is displayed in magenta. Hydrogen bond network is depicted by blue dashes. **B** Comparisons of NdhD and NdhF key proton transfer residues in NDH-1MS’, NDH-1MS (PDB 6TJV) and NDH-1L (PDB 6KHJ). Solid arrow indicates the previously proposed proton pumping pathway in NdhD4^37^. Dashed arrow indicates the putative proton pathway in NdhF4. Notice that the connection to the thylakoid lumen in NdhF4 was interrupted by non-polar residues.

Previous MD simulations of NDH-1L and NDH-1MS proposed an S-shaped proton translocation pathway connecting the N-side to the P-side via central titratable residues in each ALS^17,37^. Consistently, the N-side entrances of ALSs connect to the cytoplasm through water clusters (Fig. 5a). In NdhF4, a potential water network extends from the unknown density UNK1 in CupB towards and through the N-side channel to the central residues of NdhF4, raising the possibility that NdhF4 could take up protons directly from the UNK1-binding region (Fig. 3c, 5a). A similar hydrated route extends from the cytoplasmic side towards the central residues of NdhD4. Not all residues in these pathways fall within hydrogen-bonding distance, likely reflecting the absence of resolved water at the present resolution and the catalytic state of the complex. In NdhB, the linkage between Lys300 and Tyr236 is blocked by the conserved gating residue Phe344. This obstruction is less severe in NdhD4 and NdhF4, where the bulky Phe residues (Phe347_D4_, Phe350_F4_) can be bypassed through hydrogen bond with Ser247_D4_ and Ser255_F4_. These residues correspond to the gating region described in proton-pumping complex I homologues and may influence hydration and proton accessibility of the N-side channels^34^.

Interactions between TM7 and TM5 Lys–Glu/Asp ion pairs, together with protonation of central titratable residues, have been suggested to promote hydration and continuity of the N-side channel^38–40^. In the current structure, these ion pairs are dissociated, adopting an “open” conformation (Fig. 5a)^38,41^, and the central Lys residues (Lys252_B_, Lys256_D4_, Lys346_F4_) are predicted to be deprotonated (calculated by PropKa^32^). This configuration mirrors the closed N-side channel state of NDH-1L and complex I observed in simulations^37,38^. It is possible that PQ reduction transiently opens the N-side channel by inducing conformational changes at the gating residues, thereby enabling fuller hydration of the channel^34,38^. Although no continuous water wire connects the P-side exit (TM12 Glu/Lys) to the thylakoid lumen, structural water molecules located between TM5 and TM12 contribute to the Grotthuss-type network.

The primary distinction of the NDH-1MS’ proton translocation pathway from complex I and other NDH-1 variants lie within NdhF4. In the distal ALS of complex I and NDH-1L (NdhF1 ND5, NuoL, Nqo12), the central proton pathway connects to the P-side exit via conserved His, Lys, and Asp residues, enabling proton export^26,36^. In NdhF3, however, most key proton-transfer residues are replaced by hydrophobic residues, effectively shutting off proton translocation^17^. By contrast, NdhF4 retained most of the important titratable residues but replaced the P-side exit residues with Val338, Phe401, and Tyr402 (Fig. 5b). Impaired proton export has been demonstrated in complex I when the TM12 Lys and Asp are mutated to non-polar residues^42,43^. This critical substitution suggests that NdhF4 has no direct contact with the thylakoid lumen and is unlikely to export protons. Nevertheless, the Grotthuss-type network extending from the N-side entrance to the central pathway indicates that NdhF4 could potentially transfer protons to NdhD4 via TM5 Glu150_F4_, TM12 Glu401_D4_.

## Discussion

Here, we present the first cryo-EM structural analysis of the constitutive cyanobacterial carbon-concentrating mechanism NDH-1MS’ in both the NdhV-bound and NdhV-dissociated states. Comparison of these two states links NdhV binding to PQ destabilization, suggesting that NdhV may modulate PQ mobility. The structures also challenge the precise location of the CO_2_ hydration active site within the carbon-concentrating module. In contrast to the inducible counterpart NDH-1MS^17^, our structure showed that the previously proposed zinc-coordinating active site is instead occupied by a well-resolved phosphatidylglycerol molecule. This discrepancy in ligand assignment may reflect the improved local resolution in the present study. Recent site-directed mutagenesis of CupB residues His86, Arg91, and Glu95 in *Synechococcus elongatus* (homologous to His88, Arg93, and Glu97 in *T. vestitus*) demonstrated their essential role in carbon-concentrating activity and stable complex assembly^44,45^. In our structure, the same residues form hydrogen bonds with a PG, which may function as a “molecular bolt” reinforcing the interaction between CupB and NdhF4. This might account for the dissociation of the complex when these interactions are disrupted. Nevertheless, it remains possible that NDH-1MS and NDH-1MS’ employ distinct active sites. Although NDH-1MS’ is structurally competent to form a CO_2_ tunnel, as predicted for NDH-1MS^17^, the function of this tunnel becomes less clear if the originally proposed interface site is occupied by PG rather than a catalytic metal center. This reassignment therefore justifies the consideration of an alternative active site within the conserved internal pocket of CupB.

This pocket contains two unknown densities whose identity and function remain to be clarified. Nevertheless, their position, geometry and conserved chemical environment suggest a potential alternative active site. This possibility is supported by the strictly conserved surrounding amino acids among CupB homologues and by the geometry of the densities (Fig. 3b). Interestingly, the pocket partially resembles the zinc-less iota-carbonic anhydrase (ι-CA) identified in *Anabaena* sp. and *Bigelowiella natans*^46^. This unusual CA subclass features a metal ion-independent active site scaffolded by Tyr, Trp and Phe; however, the polar active-site cleft characteristic of ι-CA is absence in CupB. In canonical carbonic anhydrases, the metal ion lowers the pKa of substrate water, thereby enabling formation of a hydroxide nucleophile, whereas in zinc-less ι-CA this function may instead be accomplished by the active site titratable residues^47–49^. In this context, the largely hydrophobic UNK1/2 pocket in CupB could, in principle, coordinate CO_2_/HCO_3_^−^ and the substrate water through a combination of hydrophobic interactions and hydrogen bonds with the Tyr and Trp residues (Fig. 3b). The pocket may not itself function as a conventional carbonic anhydrase active site, but rather as a specialized substrate-binding environment whose catalytic competence depends on coupling to proton-pumping.

The proton transfer network connecting NdhF4 and CupB further suggests that substrate water deprotonation could be coupled to proton-pumping in the ALS, as proposed in recent mechanistic models^45,50^. However, where protons are ultimately exported remains unresolved. Based on experimentally determined structures of complex I, it has been proposed that all protons are exported in the last ALS^26,31,51^; yet NdhF4 appears structurally ill-suited for direct proton export because its P-side exit lacks the key residues. This implies that NDH-1MS’ may not follow the “domino effect” model of proton pumping^51^, or that coupling in the terminal module differs fundamentally from currently characterized homologues. Alternatively, protons could be exported from each ALS, as proposed in the “wave propagation” model^39,52^.

Given the current understanding of NDH-1^13,14,17,18,23,25,37^, we propose a tentative coupling mechanism for the NDH-1S’ module (Fig. 6), based on the “wave propagation” model^17,38–41,52–54^. We emphasize that this model applies only to NDH-1MS’ and not to NDH-1MS, as the extensive loss of proton-transfer residues in NdhF3 points to a different mechanism in the inducible complex. According to the “wave propagation” model^39^, quinone reduction and transition to the high-potential binding site stimulate sequential opening of the intra-subunit Lys–Glu/Asp ion pairs in the ALS and loading of protons into the export sites. Proton export from the distal subunit (NuoL/ND5/Nqo12) and subsequent re-protonation initial a “reverse wave” that triggers proton export in the other ALS. Provided the structural homology between NDH-1 and complex I, a similar mechanism can be hypothesized for the NDH-1 complexes. However, unlike complex I, NdhF4 cannot directly released a proton to the thylakoid lumen. Instead, the trapped positive charge in NdhF4 could promote proton export from NdhD4 which can then be re-protonated by NdhF4 via the inter-subunit Grotthuss-type network, effectively initiating the “reverse wave”. Irrespective of the exact location of the CupB active site, this step could connect NdhF4 re-protonation to uptake of a proton derived from substrate water during CO_2_ hydration. Thus, proton export from NdhD4 could create a long-range electrostatic pull that couples proton transfer from CupB to the membrane arm via NdhF4. On the other hand, under low cellular CO_2_ and high HCO_3_^−^ conditions, NdhD4 could be re-protonated directly from the cytoplasm, bypassing NdhF4 and CupB while maintaining PQ turnover and proton-pumping.

**Figure 6.**
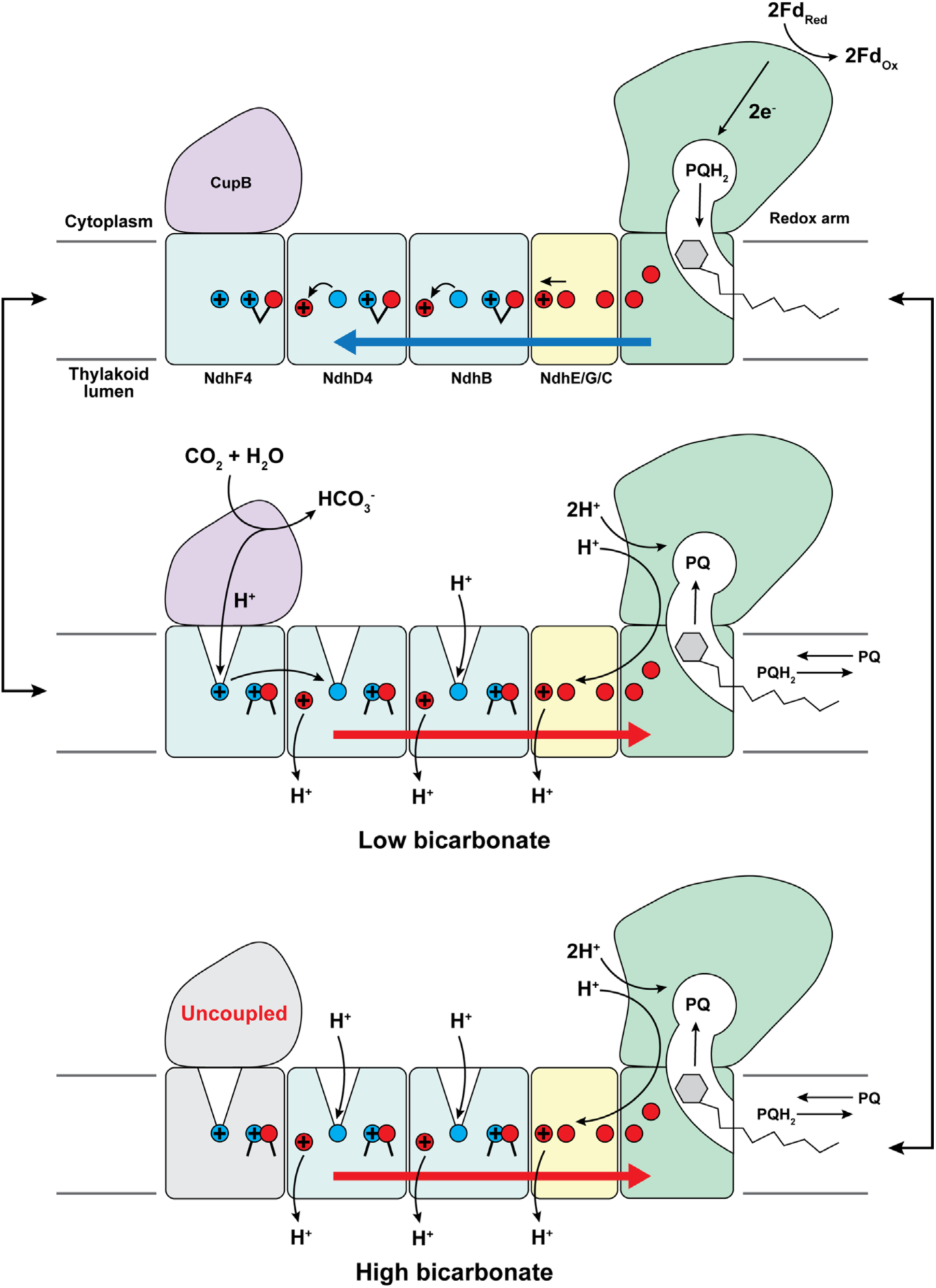
Schematic of proposed mechanism. CO_2_ hydration coupling mechanism based on the ‘wave propagation” model^39^. Red and blue circles represent Glu/Asp and Lys respectively. Protonated residues are marked by “+”. Top: Plastoquinone is reduced by two electrons from two ferredoxins. Transition of PQH_2_ from the deep binding site to the high potential binding site initiates the forward electrostatic wave (blue arrow). This sequentially opens the intra-subunit ion pair in each ALS, dehydrates N-side channel and transfers one proton to the export residue. Middle: Export of a proton from NdhD4 TM12 Glu followed by re-protonation of the central Lys generates the reverse wave (red arrow), consequently re-closes the ion pairs, opens the N-side channel and triggers proton release from NdhB and E-channel. While protons in NdhB and the E-channel are reloaded from the cytoplasm, under low bicarbonate conditions, re-protonation of NdhD4 can be coupled to CO_2_ hydration in CupB via NdhF4. Bottom: Under high bicarbonate conditions, NdhD4 could uncouple from CupB and accept a proton from the cytoplasm instead.

This working model differs from the recent “product trapping and energetic release” (PTER) model^50^ in how water deprotonation is coupled via NdhF4. The PTER model proposes that NdhF4 transiently extracts a proton from active-site-bound water and then returns this proton upon PQH2 release. By contrast, our model would entail net removal of the proton generated by substrate water deprotonation. Such a mechanism could sustain a continuous flux towards bicarbonate formation and prevent CO_2_ hydration from stalling because of proton accumulation at the catalytic site. More broadly, by employing two potentially competing proton transfer routes, NDH-1MS’ might modulate the degree of coupling between electron transfer and CO_2_ hydration. Under low bicarbonate conditions, PQ reduction could be preferentially coupled to CO_2_ hydration through the CupB-associated proton pathway. Under unusually high bicarbonate conditions, where water deprotonation becomes thermodynamically less favorable, the cytoplasmic pathway might outcompete the CupB pathway, thereby partially uncoupling carbonic anhydrase activity and lowering the effective coupling stoichiometry.

From an evolutionary perspective, a dual operating mode may have provided cyanobacteria with a selective advantage in aquatic environments, where intracellular HCO_3_^−^/CO_2_ ratios fluctuate with light intensity, pH, and nutrient status. The ability to vary the coupling between CO_2_ hydration and electron transfer would not only help maintain PQ recycling and cyclic electron flow alongside NDH-1L but also buffer the photosynthetic apparatus against sudden shifts in carbon availability. In this sense, NDH-1MS’ may represent an evolutionary compromise – a constitutive machinery that ensures baseline bicarbonate accumulation while simultaneously safeguarding photosynthetic efficiency and photoprotection across diverse ecological conditions.

## Materials and Methods

### Generation of the CupB-TS construct

Generation of the pUC18CupB-TwinStrep-tag (TS) mutant was based on the pUC18-CupS-TS plasmid that was described^17^. The *cupB* (tlr2126) gene and its corresponding upstream (US) region were amplified using genomic DNA of *Thermosynechococcus vestitus* as template. A 5’-*HindIII* and 3’-*Eco47III* restriction site was added by primer extension (primers P1-*for* and P2-*rev*; see Supplementary Table 3). The product was subcloned into pJET1.2/blunt and subsequently ligated into the pUC18-CupS-TS backbone, which was linearized by restriction with *HindIII* and *Eco47III*, exchanging *cupS* (tll0220) and its US-region. The corresponding downstream region (DS) was similarly amplified by PCR and extended with a 5’-*PstI* and a 3’-*SacI* restriction site (primers P3-*for* and P4-*rev*; see Supplementary Table 3) before being subcloned into pJET1.2/blunt and ligated into the *PstI*/*SacI* digested backbone, exchanging the *cupS* DS-region. The resulting plasmid was transformed into *T. vestitus* BP-1 WT cells by electroporation^55^. Complete segregation of the mutant allele was verified by PCR (primers P1-*for* and P4-*rev*).

### Purification of Carbon-Concentrating Complex NDH-1MS’

The CupB-TS-tag mutant was grown in a 20 L airlift photobioreactor (Bioengineering)^56^ in BG-11 liquid medium^57^ at 45 °C, with the addition of 150 µg ml^-1^ of kanamycin to uphold a selective pressure. The illumination intensity (50-200 µmol photons) was adjusted according to cell density. Mutant cultures were constantly aerated with CO_2_ (5%) before being harvested at an optical density (OD_680_) of 2. NDH-1MS**’** was isolated from the thermophilic cyanobacterium *Thermosynechococcus vestitus* BP-1 via TwinStrep(TS) affinity chromatography (IBA LifeSciences) with the TwinStrep-tag fused to the C-terminus of CupB. The solubilization of the thylakoid membranes and the purification of the complex were performed as described^14,56,58^, with the modifications as described^17^. Briefly, washed thylakoid membranes were solubilized with 1% (w/v) glyco-diosgenin (GDN) for 90 min at 20 °C. Insoluble material was removed by ultracentrifugation (45000 rpm, 1h, 4°C), and the resulting supernatant was applied to a 5 ml Strep-Tactin®XT 4Flow® high capacity column (IBA Lifesciences) for purification of the TS-tagged protein complex according to the specific workflow^14^. In this procedure, 0.02% (w/v) GDN was added to all buffers instead of 0.01% (w/v) n-dodecyl-beta-D-maltoside (DDM). After purification, the isolated protein complexes were concentrated using a spin concentrator (100 kDa cutoff) and stored at −80 °C until further analysis. Subunits composition was analyzed by mass spectrometry (see Supplementary Table 4 and Supplementary Methods).

### Cryo-EM sample preparation and data collection

To prepare cryo-EM grids, 4.5 µl protein sample (2 mg/mL) was applied to glow-discharged Quantifoil 2/1 Cu200 mesh grids, then vitrified in liquid ethane using Thermo Fisher Vitrobot Mark IV (blot force 6, blot time 6 seconds, 4 °C, 100% humidity). Sample was imaged using FEI Titan Krios operated at 300 kV, equipped with a Falcon 4i direct electron detector at the Center For Electron Microscopy in Eindhoven NanoPort Facility. Data were collected at 165,000x magnification (0.727 Å pixel size) with a defocus range of −0.8 to −1.6 µm.

### Image processing

The image processing pipeline was summarized in supplementary Figures 1. All processing steps were carried out in cryoSPARC (v4.7.0)^59^. A total of 12,384 EER movies were collected and fractionated into 50 fractions per stack without upsampling. Motion correction and contrast transfer function (CTF) estimation were performed using the Patch motion correction and Path CTF estimation jobs respectively. After discarding micrographs with CTF fit >4.0 Å, an initial set of 4000 particles were manually picked to generate a Topaz particle picking model. Three ab-initio models were reconstructed from 910,764 particles picked using the Topaz model. The relevant class (321,257 particles) was subjected to a round of heterogenous refinement into 3 classes to remove junk particles. To resolve NdhV bound particles from the NdhV dissociated state, the consensus volume was 3D classified into three classes with a mask on the NdhV binding site, enclosing NdhV/S/M and the NdhI hairpin loop. Particles of the NdhV bound state and dissociated state were polished using the Reference-based motion correction job followed by Non-uniform refinement, resulting in two consensus volumes of 2.7 Å and 2.3 Å global resolution (FSC= 0.143), respectively. To further improve the quality of the carbon-concentrating module, NdhF4 and CupB was locally refined using a focused mask to 2.4 Å global resolution.

### Model building and refinement

To construct NDH-1MS’, an initial model was assembled by rigid body fitting the NDH-1M subunits of NDH-1L (PDB 6KHI) and AlphaFold^60^ predicted models of NdhD4/F4 and CupB into the NDH-1MS’ consensus volume, followed by molecular dynamics flexible fitting (MDFF) using the Namdinator server^61^. Missing residues and ligands were then manually built in COOT^62^. This model was subjected to several rounds of iterative real-space refinement in Phenix^63^ and manual refinement in COOT. NdhF4 and CupB were extracted from NDH-1MS’ and refined on the focus refined volume using Phenix and COOT to generate the NdhF4-CupB model. A similar strategy was adopted to construct NDH-1MS’-NdhV using NDH-1MS’ as the starting model and NdhV from NDH-1L (PDB 6KHI). For structural water, the cryo-EM maps were local filtered using the Local filtering job in cryoSPARC to suppress high frequency noise. An initial set of water molecules was added to each model based on the filtered map at 2 σ using phenix.douse, then manually inspected in COOT. Only water with reasonable density fitting was retained in the final model.

## Notes

### Competing Interest Statement

The authors have declared no competing interest.

### Funding

- Deutsche Forschungsgemeinschaft: SCHU 3364/1-1, NO 836/5-1, NO836/4-1

## References

1. Field, Behrenfeld, Randerson, & Falkowski. Primary production of the biosphere: integrating terrestrial and oceanic components. Science 281, 237 true 237 237 240 true 240 240 237–40 (1998).

2. Bauwe, H., Hagemann, M. & Fernie, A. R. Photorespiration: players, partners and origin. Trends Plant Sci 15, 330 true 330 330 336 true 336 336 330–6 (2010).

3. Tcherkez, G. The mechanism of Rubisco-catalysed oxygenation. Plant Cell Env. 39, 983 true 983 983 997 true 997 997 983–97 (2015).

4. Smith, E. N., van Aalst, M., Weber, A. P. M., Ebenhöh, O. & Heinemann, M. Alternatives to photorespiration: A system-level analysis reveals mechanisms of enhanced plant productivity. Sci Adv 11, Omit eadt9287 eadt9287 eadt9287 (2025).

5. Rae, B. D., Long, B. M., Badger, M. R. & Price, G. D. Functions, compositions, and evolution of the two types of carboxysomes: polyhedral microcompartments that facilitate CO2 fixation in cyanobacteria and some proteobacteria. Microbiol Mol Biol Rev 77, 357 true 357 357 379 true 379 379 357–79 (2013).

6. Kupriyanova, E. V., Pronina, N. A. & Los, D. A. Adapting from Low to High: An Update to CO2-Concentrating Mechanisms of Cyanobacteria and Microalgae. Plants Basel 12, (2023).

7. Maberly, S. C. & Gontero, B. Ecological imperatives for aquatic CO2-concentrating mechanisms. J Exp Bot 68, 3797 true 3797 3797 3814 true 3814 3814 3797–3814 (2017).

8. Omata, T., Takahashi, Y., Yamaguchi, O. & Nishimura, T. Structure, function and regulation of the cyanobacterial high-affinity bicarbonate transporter, BCT1. Funct Plant Biol 29, 151 true 151 151 159 true 159 159 151–159 (2002).

9. Price, G. D., Shelden, M. C. & Howitt, S. M. Membrane topology of the cyanobacterial bicarbonate transporter, SbtA, and identification of potential regulatory loops. Mol Membr Biol 28, 265 true 265 265 275 true 275 275 265–75 (2011).

10. Ohkawa, H., Pakrasi, H. B. & Ogawa, T. Two types of functionally distinct NAD(P)H dehydrogenases in Synechocystis sp. strain PCC6803. J Biol Chem 275, 31630 true 31630 31630 31634 true 31634 31634 31630–4 (2000).

11. Maeda, S.-I., Badger, M. R. & Price, G. D. Novel gene products associated with NdhD3/D4-containing NDH-1 complexes are involved in photosynthetic CO2 hydration in the cyanobacterium, Synechococcus sp. PCC7942. Mol Microbiol 43, 425 true 425 425 435 true 435 435 425–35 (2002).

12. Price, G. D., Maeda, S.-I., Omata, T. & Badger, M. R. Modes of active inorganic carbon uptake in the cyanobacterium, Synechococcus sp. PCC7942. Funct Plant Biol 29, 131 true 131 131 149 true 149 149 131–149 (2002).

13. Laughlin, T. G., Savage, D. F. & Davies, K. M. Recent advances on the structure and function of NDH-1: The complex I of oxygenic photosynthesis. Biochim Biophys Acta Bioenerg 1861, 148254 true 148254 148254 148254 (2020).

14. Schuller, J. M. et al. Structural adaptations of photosynthetic complex I enable ferredoxin-dependent electron transfer. Science 363, 257 true 257 257 260 true 260 260 257–260 (2018).

15. Kramer, D. M. & Evans, J. R. The Importance of Energy Balance in Improving Photosynthetic Productivity. Plant Physiol. 155, 70–78 (2011).

16. Yu, H., Schut, G. J., Haja, D. K., Adams, M. W. W. & Li, H. Evolution of complex I-like respiratory complexes. J Biol Chem 296, 100740 true 100740 100740 100740 (2021).

17. Jan M. Schuller et al. Redox-coupled proton pumping drives carbon concentration in the photosynthetic complex I. Nat Commun 11, 494 true 494 494 494 (2020).

18. Battchikova, N., Eisenhut, M. & Aro, E.-M. Cyanobacterial NDH-1 complexes: novel insights and remaining puzzles. Biochim Biophys Acta 1807, 935 true 935 935 944 true 944 944 935–44 (2010).

19. Shibata, M. et al. Distinct constitutive and low-CO2-induced CO2 uptake systems in cyanobacteria: genes involved and their phylogenetic relationship with homologous genes in other organisms. Proc Natl Acad Sci U A 98, 11789 true 11789 11789 11794 true 11794 11794 11789–94 (2001).

20. Ohkawa, H., Price, G. D., Badger, M. R. & Ogawa, T. Mutation of ndh genes leads to inhibition of CO(2) uptake rather than HCO(3)(-) uptake in Synechocystis sp. strain PCC 6803. J. Bacteriol. 182, 2591 true 2591 2591 2596 true 2596 2596 2591–6 (2000).

21. Klughammer, B., Sültemeyer, D., Badger, M. R. & Price, G. D. The involvement of NAD(P)H dehydrogenase subunits, NdhD3 and NdhF3, in high-affinity CO2 uptake in Synechococcus sp. PCC7002 gives evidence for multiple NDH-1 complexes with specific roles in cyanobacteria. Mol Microbiol 32, 1305 true 1305 1305 1315 true 1315 1315 1305–15 (1999).

22. Arteni, A. A. et al. Structural characterization of NDH-1 complexes of Thermosynechococcus elongatus by single particle electron microscopy. Biochim Biophys Acta 1757, 1469 true 1469 1469 1475 true 1475 1475 1469–75 (2006).

23. Pan, X. et al. Structural basis for electron transport mechanism of complex I-like photosynthetic NAD(P)H dehydrogenase. Nat Commun 11, 610 true 610 610 610 (2020).

24. Laughlin, T. G., Bayne, A. N., Trempe, J.-F., Savage, D. F. & Davies, K. M. Structure of the complex I-like molecule NDH of oxygenic photosynthesis. Nature 566, 411 true 411 411 414 true 414 414 411–414 (2019).

25. Zhang, C. et al. Structural insights into NDH-1 mediated cyclic electron transfer. Nat Commun 11, 888 true 888 888 888 (2020).

26. Kravchuk, V. et al. A universal coupling mechanism of respiratory complex I. Nature 609, 808 true 808 808 814 true 814 814 808–814 (2022).

27. Chovancova, E. et al. CAVER 3.0: a tool for the analysis of transport pathways in dynamic protein structures. PLoS Comput Biol 8, Omit e1002708 e1002708 e1002708 (2012).

28. Chung, I., Grba, D. N., Wright, J. J. & Hirst, J. Making the leap from structure to mechanism: are the open states of mammalian complex I identified by cryoEM resting states or catalytic intermediates? Curr. Opin. Struct. Biol. 77, 102447 (2022).

29. Zhu, J., Vinothkumar, K. R. & Hirst, J. Structure of mammalian respiratory complex I. Nature 536, 354 true 354 354 358 true 358 358 354–358 (2016).

30. Chung, I. et al. Cryo-EM structures define ubiquinone-10 binding to mitochondrial complex I and conformational transitions accompanying Q-site occupancy. Nat Commun 13, 2758 true 2758 2758 2758 (2022).

31. Kampjut, D. & Sazanov, L. A. The coupling mechanism of mammalian respiratory complex I. Science 370, (2020).

32. Olsson, M. H. M., Søndergaard, C. R., Rostkowski, M. & Jensen, J. H. PROPKA3: Consistent Treatment of Internal and Surface Residues in Empirical pKa Predictions. J Chem Theory Comput 7, 525 true 525 525 537 true 537 537 525–37 (2011).

33. Ivanov, B. S., Bridges, H. R., Jarman, O. D. & Hirst, J. Structure of the turnover-ready state of an ancestral respiratory complex I. Nat Commun 15, 9340 true 9340 9340 9340 (2024).

34. Parey, K. et al. High-resolution structure and dynamics of mitochondrial complex I-Insights into the proton pumping mechanism. Sci Adv 7, Omit eabj3221 eabj3221 eabj3221 (2021).

35. Grba, D. N. & Hirst, J. Mitochondrial complex I structure reveals ordered water molecules for catalysis and proton translocation. Nat Struct Mol Biol 27, 892 true 892 892 900 true 900 900 892–900 (2020).

36. Grba, D. N., Chung, I., Bridges, H. R., Agip, A.-N. A. & Hirst, J. Investigation of hydrated channels and proton pathways in a high-resolution cryo-EM structure of mammalian complex I. Sci Adv 9, Omit eadi1359 eadi1359 eadi1359 (2023).

37. Saura, P. & Kaila, V. R. I. Molecular dynamics and structural models of the cyanobacterial NDH-1 complex. Biochim Biophys Acta Bioenerg 1860, 201 true 201 201 208 true 208 208 201–208 (2018).

38. Di Luca, A., Gamiz-Hernandez, A. P. & Kaila, V. R. I. Symmetry-related proton transfer pathways in respiratory complex I. Proc Natl Acad Sci U A 114, Omit E6314–E6321 (2017).

39. Kaila, V. R. I. Long-range proton-coupled electron transfer in biological energy conversion: towards mechanistic understanding of respiratory complex I. J R Soc Interface 15, (2018).

40. Röpke, M., Saura, P., Riepl, D., Pöverlein, M. C. & Kaila, V. R. I. Functional Water Wires Catalyze Long-Range Proton Pumping in the Mammalian Respiratory Complex I. J Am Chem Soc 142, 21758 true 21758 21758 21766 true 21766 21766 21758–21766 (2020).

41. Mühlbauer, M. E. et al. Water-Gated Proton Transfer Dynamics in Respiratory Complex I. J. Am. Chem. Soc. 142, 13718 true 13718 13718 13728 true 13728 13728 13718–13728 (2020).

42. Beghiah, A. et al. Dissected antiporter modules establish minimal proton-conduction elements of the respiratory complex I. Nat Commun 15, 9098 true 9098 9098 9098 (2024).

43. Nakamaru-Ogiso, E. et al. The Membrane Subunit NuoL(ND5) Is Involved in the Indirect Proton Pumping Mechanism of Escherichia coli Complex I. J Biol Chem 285, 39070 true 39070 39070 39078 true 39078 39078 39070–39078 (2010).

44. Artier, J. et al. Modeling and mutagenesis of amino acid residues critical for CO2 hydration by specialized NDH-1 complexes in cyanobacteria. Biochim Biophys Acta Bioenerg 1863, 148503 true 148503 148503 148503 (2021).

45. Walker, R. M., Zhang, M. & Burnap, R. L. Elucidating the role of primary and secondary sphere Zn2+ ligands in the cyanobacterial CO2 uptake complex NDH-14: The essentiality of arginine in zinc coordination and catalysis. Biochim Biophys Acta Bioenerg 1865, 149149 true 149149 149149 149149 (2024).

46. Hirakawa, Y. et al. Characterization of a novel type of carbonic anhydrase that acts without metal cofactors. BMC Biol 19, 105 true 105 105 105 (2021).

47. Boone, C. D., Pinard, M., McKenna, R. & Silverman, D. Catalytic mechanism of α-class carbonic anhydrases: CO2 hydration and proton transfer. Subcell Biochem 75, 31 true 31 31 52 true 52 52 31–52 (2014).

48. Rowlett, R. S. Structure and catalytic mechanism of the beta-carbonic anhydrases. Biochim Biophys Acta 1804, 362 true 362 362 373 true 373 373 362–73 (2009).

49. Lindskog, S. Structure and mechanism of carbonic anhydrase. Pharmacol Ther 74, 1 true 1 1 20 true 20 20 1–20 (1997).

50. Zhang, Z., Zhang, M. & Burnap, R. L. Action at a distance: The remarkable coupling of CO2 uptake to electron transfer in specialized cyanobacterial NDH-1 complexes. Proc Natl Acad Sci USA 122, Omit e2511786122 e2511786122 e2511786122 (2025).

51. Sazanov, L. A. From the ‘black box’ to ‘domino effect’ mechanism: what have we learned from the structures of respiratory complex I. Biochem J 480, 319 true 319 319 333 true 333 333 319–333 (2023).

52. Kaila, V. R. I. Resolving Chemical Dynamics in Biological Energy Conversion: Long-Range Proton-Coupled Electron Transfer in Respiratory Complex I. Acc Chem Res 54, 4462 true 4462 4462 4473 true 4473 4473 4462–4473 (2021).

53. Kaila, V. R. I., Wikström, M. & Hummer, G. Electrostatics, hydration, and proton transfer dynamics in the membrane domain of respiratory complex I. Proc Natl Acad Sci U A 111, 6988 true 6988 6988 6993 true 6993 6993 6988–93 (2014).

54. Mühlbauer, M. E., Gamiz-Hernandez, A. P. & Kaila, V. R. I. Functional Dynamics of an Ancient Membrane-Bound Hydrogenase. J. Am. Chem. Soc. 143, 20873 true 20873 20873 20883 true 20883 20883 20873–20883 (2021).

55. Iwai, M., Katoh, H., Katayama, M. & Ikeuchi, M. Improved genetic transformation of the thermophilic cyanobacterium, Thermosynechococcus elongatus BP-1. Plant Cell Physiol 45, 171 true 171 171 175 true 175 175 171–5 (2004).

56. Wulfhorst, H., Franken, L. E., Wessinghage, T., Boekema, E. J. & Nowaczyk, M. M. The 5 kDa protein NdhP is essential for stable NDH-1L assembly in Thermosynechococcus elongatus. PLoS One 9, Omit e103584 e103584 e103584 (2014).

57. Stanier, R. Y., Deruelles, J., Rippka, R., Herdman, M. & Waterbury, J. B. Generic Assignments, Strain Histories and Properties of Pure Cultures of Cyanobacteria. Microbiology 111, 1 true 1 1 61 true 61 61 1–61 (1979).

58. Kuhl, H. et al. Towards structural determination of the water-splitting enzyme. Purification, crystallization, and preliminary crystallographic studies of photosystem II from a thermophilic cyanobacterium. J Biol Chem 275, 20652 true 20652 20652 20659 true 20659 20659 20652–9 (2000).

59. Punjani, A., Rubinstein, J. L., Fleet, D. J. & Brubaker, M. A. cryoSPARC: algorithms for rapid unsupervised cryo-EM structure determination. Nat Methods 14, 290 true 290 290 296 true 296 296 290–296 (2017).

60. Jumper, J. et al. Highly accurate protein structure prediction with AlphaFold. Nature 596, 583 true 583 583 589 true 589 589 583–589 (2021).

61. Kidmose, R. T. et al. Namdinator - automatic molecular dynamics flexible fitting of structural models into cryo-EM and crystallography experimental maps. IUCrJ 6, 526 true 526 526 531 true 531 531 526–531 (2019).

62. Emsley, P., Lohkamp, B., Scott, W. G. & Cowtan, K. Features and development of Coot. Acta Crystallogr Biol Crystallogr 66, 486 true 486 486 501 true 501 501 486–501 (2010).

63. Liebschner, D. et al. Macromolecular structure determination using X-rays, neutrons and electrons: recent developments in Phenix. Acta Crystallogr Struct Biol 75, 861 true 861 861 877 true 877 877 861–877 (2019).

64. Ashkenazy, H. et al. ConSurf 2016: an improved methodology to estimate and visualize evolutionary conservation in macromolecules. Nucleic Acids Res 44, 50 Omit W344–50 (2016).

65. Tian, W., Chen, C., Lei, X., Zhao, J. & Liang, J. CASTp 3.0: computed atlas of surface topography of proteins. Nucleic Acids Res. 46, W363–W367 (2018).

