## Supplementary Information for "Structural Basis of Redox-Coupled CO_2_ hydration by the Cyanobacterial NDH-1MS’ complex"

**Supplementary Methods**
**Supplementary Tables 1 - 4**
**Supplementary Figures 1 - 8**

### Supplementary Methods

#### Mass Spectrometric analysis

Masses of small intact proteins of NDH-1MS' were determined by matrix assisted laser desorption/ionic time-of-flight (ToF) tandem mass spectrometry (MS) as described[1]. Isolated NDH-1MS' was mixed in 1:1 and 1:5 ratios with saturated matrix solution (sinapic acid in 1 % (v/v) trifluoroacetic acid and 60 % (v/v) acetonitrile) before 1 µl of the mixture was spread and air-dried on the target plate as duplicates. The measurement was performed on an Ultraflex III (Bruker, Daltonics). The sample was ionized by a pulsed ultraviolet laser (100 µJ at 337 nm) at a repetition rate of 50 Hz before being further accelerated (19 kV). Masses were determined by the time-of-flight in high vacuum at $8x10^{-7}$ mbar via micro channel plate detectors. Intact protein mass was determined in the m/z range of 3 to 20 kDa. Spectra analysis was performed using the open-source software mMass (version 5.5.0 2013) as a sum of 10-50 laser shots[2].

Liquid Chromatography-Nanospray Ionization-Tandem mass spectrometry (LC-NSI-MS/MS) was performed at an Orbitrap Elite mass spectrometer (Thermo Fisher Scientific) using the Proteome Discoverer software (version 2.3). Sample preparation, setup and analysis were performed according to previous studies[3,4] with minor changes. Briefly, in-gel digestion and rehydration (0.125 µg trypsin, 25 mM $NH_4HCO_3$, 10 % (v/v) MeOH) were carried out at 37 °C overnight following dehydration in 100 % (v/v) acetonitrile for 5 min and drying using a rotary evaporator (Concentrator Plus, Eppendorf). Peptides were extracted by incubation in extraction buffer (50 % (v/v) acetonitrile and 1 % (v/v) formic acid) for 20 min. The extracted peptides were dried again before being resuspended in solution A (0.1% (v/v) formic acid, 2% (v/v) acetonitrile) and centrifuged (1 min, RT, 13200 rpm, Eppendorf, SL 073) for the removal of precipitated molecules.

Peptides were subsequently applied on a nanoACQUITY UPLC Symmetry C18 Trap column (Waters GmbH) followed by a ACQUITY UPLC BEH C18 column (Waters, GmbH). Elution was performed with a flow rate of 0.4 µl/min at 55 °C in a discontinuous gradient of solution A to solution B (0.1% (v/v) formic acid in acetonitrile) over 60 min as described[3]. The eluted peptides were transferred via SilicaTip emitter (30 µm) for nanospray ionization at 1.5 -1.8 kV. The mass spectra were recorded at a range of 300 – 2,000 m/z with a resolution of 240,000. Tandem MS (MS/MS) spectra of the 20 intense precursor ions in the ion trap were recorded (Top 20 method) and the collision-induced dissociation (CID) was set to 35%. The dynamic exclusion of ions was activated with a one-minute time window and a repetition count of 1. Single charged ions and ions with an unassigned charge state were rejected from the spectra. Analysis was performed using a modified database from *T. vestitus* BP-1 (Proteom-ID: UP000000440)[5] to which common contaminants as human keratin from the Global Proteom Machine database were. Furthermore, sequences of the tagged proteins and posttranslational modifications (PTMs) as N-terminal acetylation (NTA) and oxidation of methio-nine residues were added. The False Discovery Rate (FDR) was set to 1% and the assignment of the spectra was conducted with the Sequest-algorithm[6] automatically by the Proteom Discoverer software.

### Supplementary tables and figures

**Supplementary Table 1. Cryo-EM data collection, refinement and validation statistics.**

| | NDH-1MS‘ (EMDB-59241) (PDB 32XI) | NDH-1MS‘-NdhV (EMDB- 59242) (PDB 32XJ) | NDH-1MS‘ (focused refinement) (EMDB- 59243) (PDB 32XK) |
|---|---|---|---|
| Data collection and processing | | | |
| Magnification | | 165,000 | |
| Voltage (kV) | | 300 | |
| Electron exposure (e–/Å$^2$) | | 50 | |
| Defocus range (µm) | | -0.8 to -1.6 | |
| Pixel size (Å) | | 0.727 | |
| Symmetry imposed | | C1 | |
| Initial particle images (no.) | | 910,794 | |
| Final particle images (no.) | 112,985 | 52,063 | 112,985 |
| Map resolution (Å) | 2.34 | 2.67 | 2.44 |
| FSC threshold | 0.143 | 0.143 | 0.143 |
| Map resolution range (Å) | 2.3 – 3.5 | 2.5 – 4.0 | 2.3 – 3.2 |
| Refinement | | | |
| Model resolution (Å) | 2.34 | 2.67 | 2.44 |
| FSC threshold | 0.143 | 0.143 | 0.143 |
| Map sharpening *B* factor (Å$^2$) | -42.7 | -41.9 | -52.8 |
| Model composition | | | |
| Non-hydrogen atoms | 33,984 | 34,731 | 7,709 |
| Protein residues | 4,113 | 4,214 | 914 |
| Ligands | 35 | 36 | 11 |
| *B* factors (Å$^2$) | | | |
| Protein (mean) | 58.93 | 60.99 | 52.10 |
| Ligand (mean) | 78.21 | 81.47 | 72.03 |
| R.m.s. deviations | | | |
| Bond lengths (Å) | 0.00 | 0.00 | 0.00 |
| Bond angles (°) | 0.02 | 0.02 | 0.04 |
| Validation | | | |
| MolProbity score | 1.54 | 1.60 | 1.30 |
| Clashscore | 6.35 | 8.03 | 5.19 |
| Poor rotamers (%) | 0.44 | 0.26 | 0.13 |
| Ramachandran plot | | | |
| Favored (%) | 96.88 | 97.1 | 97.91 |
| Allowed (%) | 3.09 | 2.8 | 2.09 |
| Disallowed (%) | 0.02 | 0.1 | 0.00 |

**Supplementary Table 2. NDH-1S/S' subunit interactions**

**Side chain interactions between NdhD4-NdhF4 and NdhD3-NdhF3**

| NdhF4 | Distance (Å) | NdhD4 | NdhF3 | Distance (Å) | NdhD3 |
|---|---|---|---|---|---|
| GLN 75 [OE1] | 3.30 | ASN 475 [ND2] | | -- | |
| TYR 157 [OH] | 3.86 | ASN 435 [ND2] | TYR 157 [OH] | 2.20 | ASN 431 [OD1] |
| | -- | | ASN 165 [OD1] | 3.78 | ASN 366 [ND2] |
| ARG 181 [NE] | 3.07 | SER 427 [OG] | | -- | |
| ASP 556 [OD2] | 3.60 | HIS 232 [NE2] | ASP 557 [OD2] | 2.36 | HIS 228 [NE2] |
| ASP 556 [OD2] | 3.48 | TYR 308 [OH] | ASP 557 [OD2] | 2.91 | TYR 287 [OH] |
| ASP 556 [OD1] | 3.55 | HIS 232 [NE2] | | -- | |
| ASP 561 [OD1] | 2.81 | LYS 168 [NZ] | ASP 562 [OD2] | 2.67 | LYS 164 [NZ] |
| ASN 565 [OD1] | 3.12 | TYR 164 [OH] | | -- | |
| ASN 565 [OD1] | 3.65 | LYS 168 [NZ] | | -- | |

**Side chain interactions between CupB-NdhF4 and CupA-NdhF3**

| CupB | Distance (Å) | NdhF4 | CupA | Distance (Å) | NdhF3 |
|---|---|---|---|---|---|
| ASP 92 [OD1] | 3.06 | ARG 33 [NH1] | | -- | |
| ASP 92 [OD2] | 3.01 | ARG 33 [NH2] | | -- | |
| GLU 97 [OE2] | 3.91 | ARG 37 [NH2] | | -- | |
| | -- | | ARG 146 [NH2] | 3.55 | GLU 114 [OE2] |
| ASP 115 [OD1] | 2.99 | ARG 447 [NH1] | | -- | |
| | -- | | ASP 158 [OD1] | 3.93 | ARG 448 [NE] |
| | -- | | ASP 158 [OD2] | 3.27 | ARG 448 [NH1] |
| ASP 115 [OD1] | 2.92 | ARG 448 [NH2] | ASP 158 [OD1] | 3.62 | ARG 448 [NH2] |
| ASP 119 [OD1] | 3.07 | ARG 448 [NH2] | ASP 162 [OD1] | 2.72 | ARG 448 [NH2] |
| | -- | | GLU 197 [OE1] | 2.68 | ARG 374 [NE] |
| | -- | | ARG 295 [NH2] | 2.95 | TYR 112 [OH] |
| ASN 258 [ND2] | 3.67 | GLU 249 [OE2] | | -- | |
| GLN 260 [NE2] | 3.38 | GLN 305 [OE1] | GLN 303 [NE2] | 3.68 | GLN 305 [OE1] |
| ARG 266 [NH2] | 3.02 | THR 366 [OG1] | | -- | |
| ARG 266 [NH2] | 3.99 | GLU 367 [OE1] | | -- | |
| ARG 266 [NH1] | 3.73 | GLU 367 [OE2] | | -- | |
| ASP 317 [OD1] | 3.56 | ARG 32 [NH2] | | -- | |

Polar interacting pairs were calculated using the PISA server. For NDH-1S interactions, PDB 6TJV was used for the analysis. Conserved interactions are highlighted in green.

**Supplementary Table 3. List of primers used for the construction of the *T. vestitus* pUC18CupB-TS**

| **Primer Name** | **DNA Sequence (5' - 3')** | **Function** |
|---|---|---|
| P1-*for* | AAGCTTGCGTGGCTACTGCTGTAATG | Amplification of cupB (tlr2126) + US region; introduction of *HindIII* and *Eco47III* restriction sites, P1-*for* used for Segregation check |
| P2-*rev* | AGCGCTGCTAAGTTGAACATTCCAGA | |
| P3-*for* | CTGCAGGGGACTCTTTTAGCCCTTGA | Amplification of cupB DS region; introduction of *PstI* and SacI; P4-*rev* used for Segregation check |
| P4-*rev* | GAGCTCACAACGGCAATGGCAGCAAG | |

US = Upstream, DS = Downstream

**Supplementary Table 4. Tandem-LC-MS analysis of NDH-1MS'**

| Subunit[a] | ORF | kDa[b] | TMH[c] | XC[d] | Coverage[e] |
|---|---|---|---|---|---|
| NdhA | tlr0667 | 41.3 | 8 | 74.68 | 25.33 |
| NdhB | tll0045 | 55.1 | 14 | 52.61 | 7.51 |
| NdhC | tlr1429 | 13.7 | 3 | 3.89 | 15.91 |
| NdhD4 | tlr2125 | 53.4 | 14 | 11.91 | 4.82 |
| NdhE | tlr0670 | 11.2 | 3 | 16.79 | 14.85 |
| NdhF3 | tlr0904 | 66.3 | 16 | 2.62 | 3.27 |
| NdhF4 | tlr2124 | 66.1 | 16 | 114.40 | 11 |
| NdhG | tlr0669 | 21.6 | 5 | 17.79 | 25.5 |
| NdhH | tlr1288 | 45.2 | - | 468.89 | 51.02 |
| NdhI | tlr0668 | 22.4 | - | 157.38 | 47.69 |
| NdhJ | tlr1430 | 19.2 | - | 145.76 | 71.43 |
| NdhK | tlr0705 | 25.7 | - | 170.04 | 41.77 |
| NdhL | tsr0706 | 11.6 | 2 | 5.44 | 11.84 |
| NdhM | tll0447 | 12.6 | - | 161.86 | 49.55 |
| NdhN | tlr1130 | 8.3 | - | 104.65 | 61.33 |
| NdhO | tsl0017 | 7.9 | - | 22.7 | 67.14 |
| NdhS | tlr0636 | 8.1 | - | 19.45 | 45.95 |
| NdhV | tlr0472 | 13.6 | - | 11.27 | 19.2 |
| CupA | tlr0906 | 50.9 | - | 7.48 | 8.42 |
| CupB-TS | tlr2126 | 46.2 | - | 266.98 | 63.66 |

[a] results were filtered for known NDH-1MS' subunits

[b] calculated molecular weight

[c] number of transmembrane helices

[d] sequest protein score

[e] sequence coverage (percent)

**A** **B**

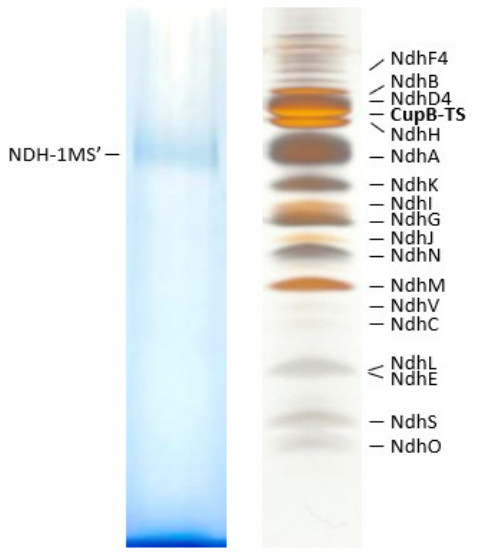

**C**

| Subunit | Calculated average mass [kDa] | Measured average mass [kDa] | Delta mass [Da] | Charge | Modification[a] |
|---|---|---|---|---|---|
| NdhJ | 19.212 | 19.216 | 4 | +1 | minus M1 |
| NdhN | 16.505 | 16.503 | 2 | +1 | minus M1 |
| NdhV | 13.600 | 13.597 | 3 | +1 | M22 (minus M1) |
| NdhM | 12.567 | 12.567 | 0 | +1 | - |
| NdhE | 11.161 | 11.161 | 0 | +1 | Formyl-M1 |
| NdhJ | 9.606 | 9.603 | 3 | +2 | minus M1 |
| NdhL | 8.559 | 8.600 | 1 | +1 | Formyl-M1 |
| NdhN | 8.251 | 8.255 | 4 | +2 | - |
| NdhS | 8.083 | 8.083 | 0 | +1 | M37 (minus M1) |
| NdhO | 7.736 | 7.735 | 1 | +1 | minus M1 |
| NdhM | 6.284 | 6.286 | 2 | +2 | - |

[a] predicted common modification based on the measured mass and/or corrected first N-terminal amino acid residue

**D**

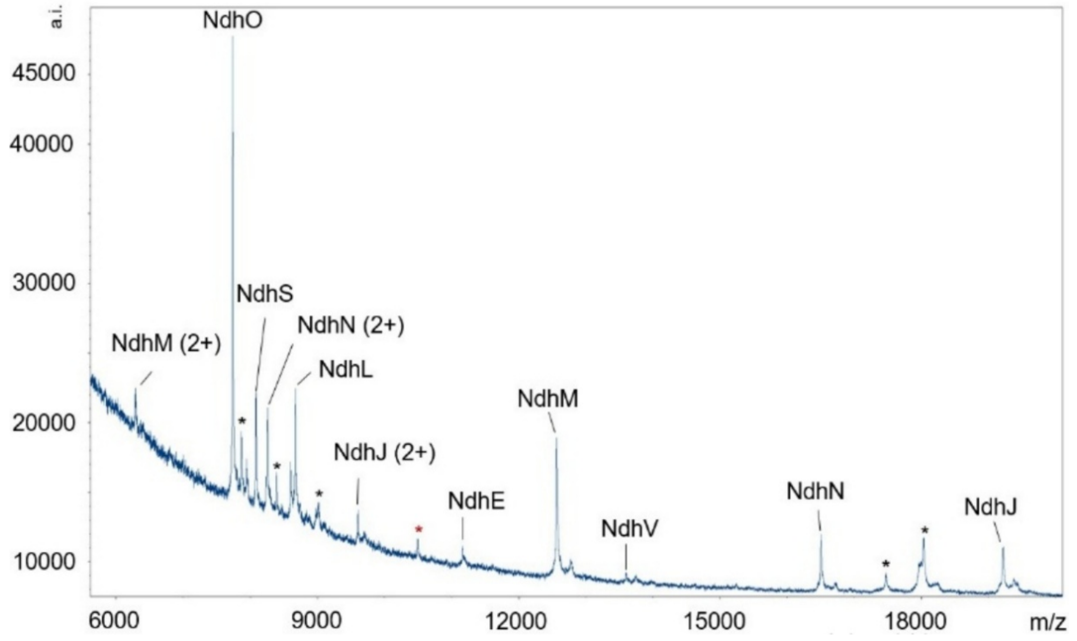

**Supplementary figure 1. Mass spectrometry analysis and verification of subunit composition of NDH-1MS’ from *Thermosynechococcus vestitus*.** **A** Blue-Native PAGE[7] and SDS-PAGE[8] (**B**) were performed to verify the integrity of the purified protein complex that was subsequently analysed by **(C/D)** Maldi-ToF mass spectrometry was performed in a *m/z* range of 5.8-20 kDa. NdhJ, NdhM and NdhN are present as single and double charged ($2^+$) ions. Red asterisk indicates the mass of a putative protein that could not be detected via tandem LC-MS, whereas the peaks labelled with black asterisks were identified as masses corresponding to phycobiliproteins (data not shown).

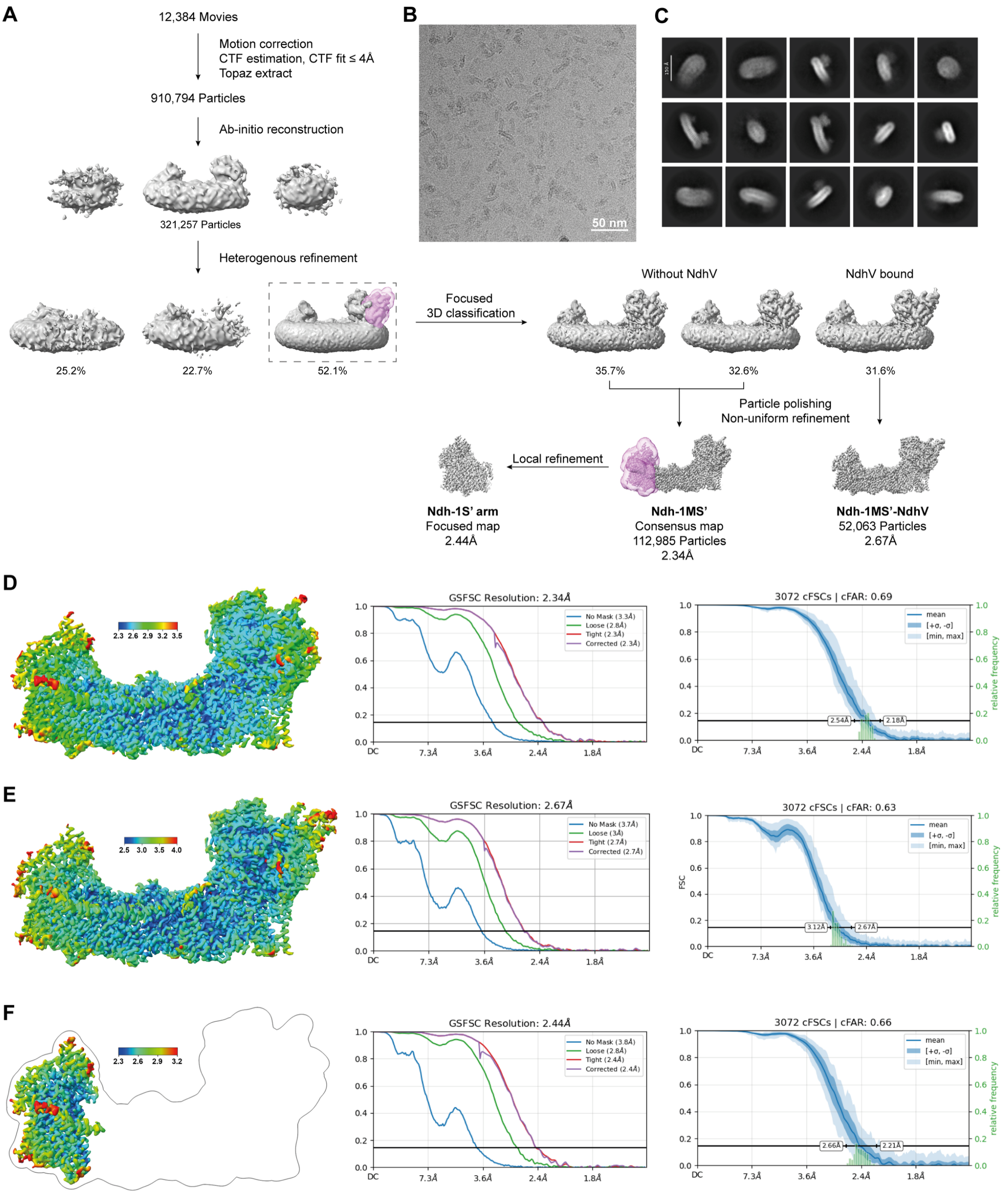

**Supplementary figure 2. Cryo-EM data processing. A** Summary of the cryo-EM data processing pipeline. **B** Representative micrograph collected at 165K magnification. **C** 2D Classes of the initial reconstruction. Local (left) and global (right) resolution of **D** NDH-1MS', **E** NDH-1MS'-NdhV and **F** focus refinement of NDH-1MS'.

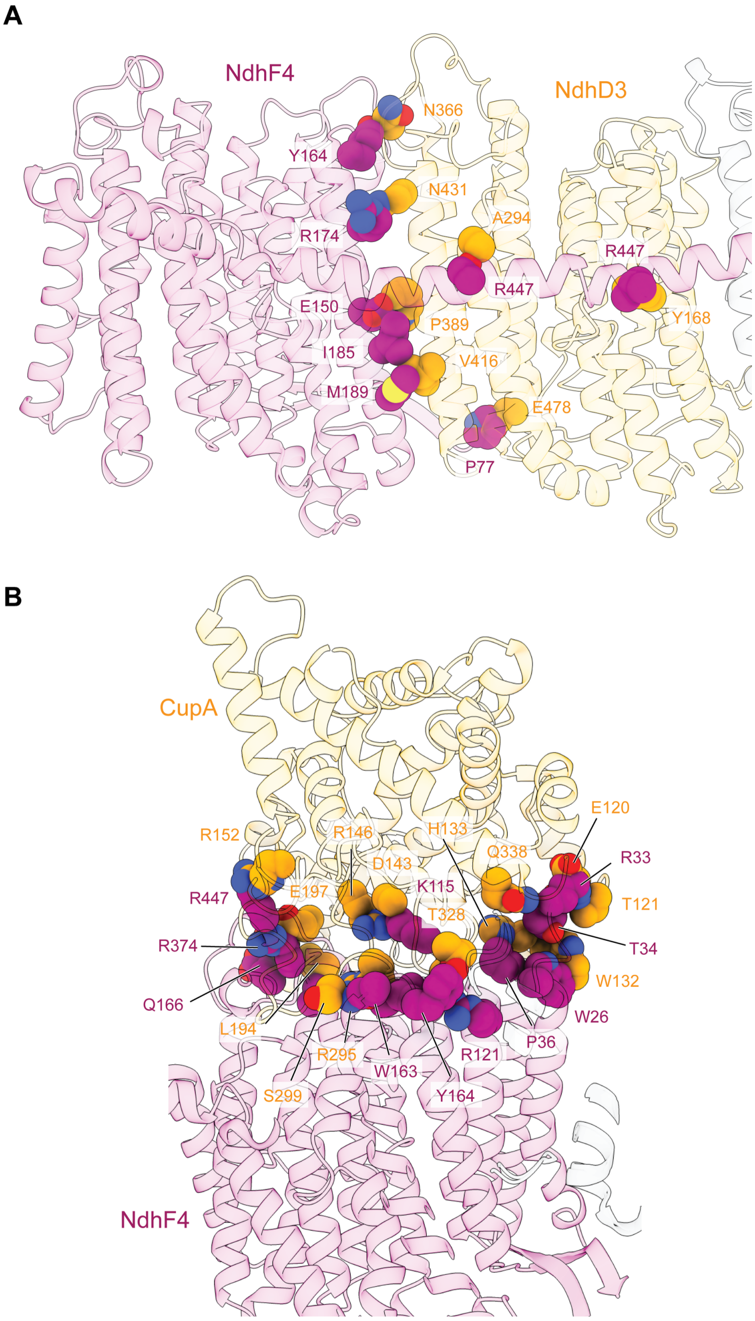

**Supplementary figure 3. Steric hindrance forbids assembly of NDH-1S/S' chimera.** NdhD3 and CupA were aligned on NdhD4 and CupB respectively. Clashing residues at **A** NdhF4-NdhD3 interface and **B** NdhF4-CupA interface are shown in sphere. Clash was defined as overlapping Van der Waals radius of ≥ 0.6 Å.

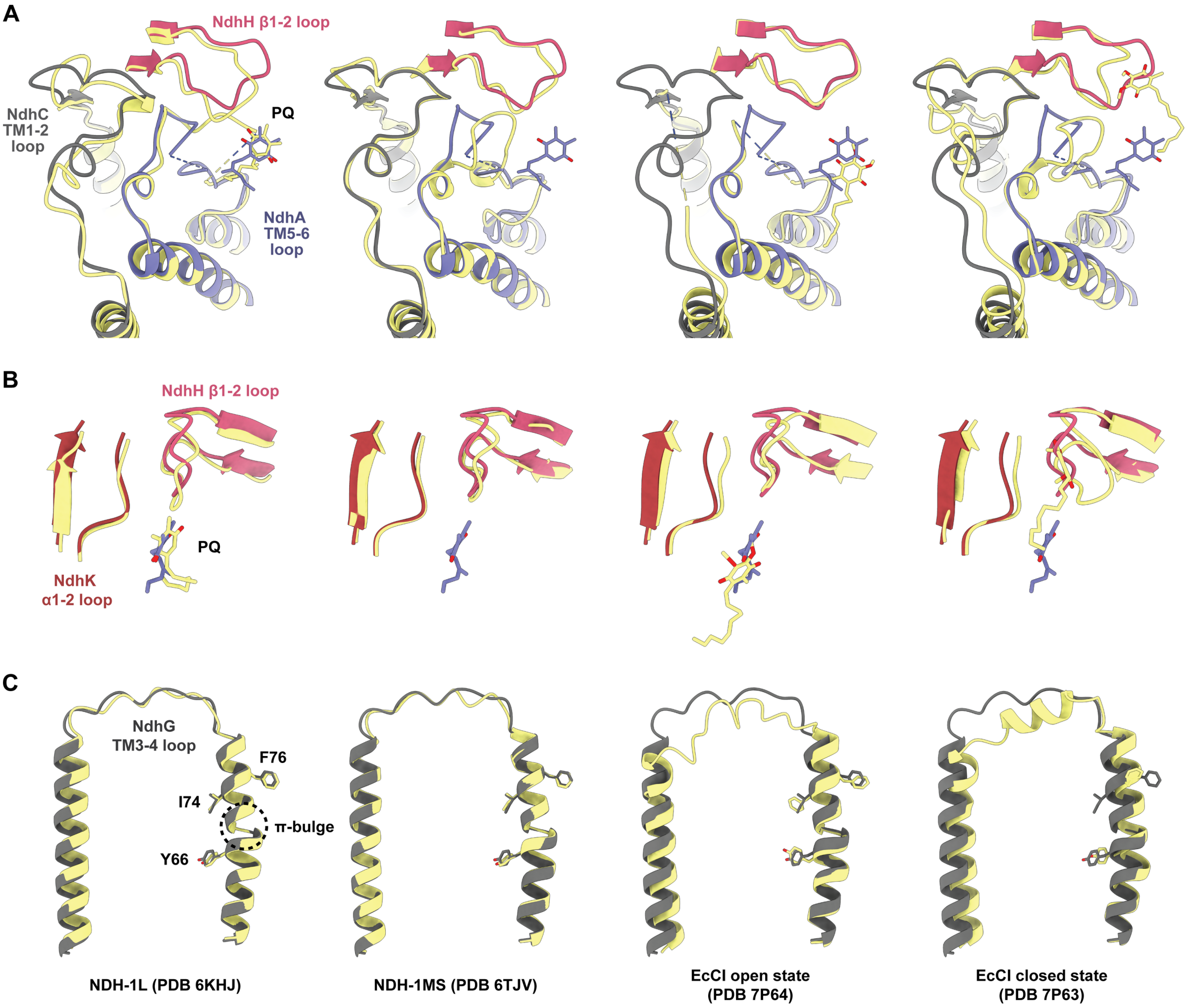

**Supplementary figure 4. Comparison of key structures in NDH-1MS' with NDH-1L, NDH-1MS and respiratory complex I. A** Comparison of NdhA/C/H (NuoH/A/CD) loops and quinone binding position. **B** Comparison of NdhK (NuoB) loop. **C** Comparison of NdhG (NuoJ) loop. **A-C** NDH-1MS' is colored as in Fig. 1. Other NDH-1 and complex I structures are colored in yellow. EcCI: *E. coli* complex I.

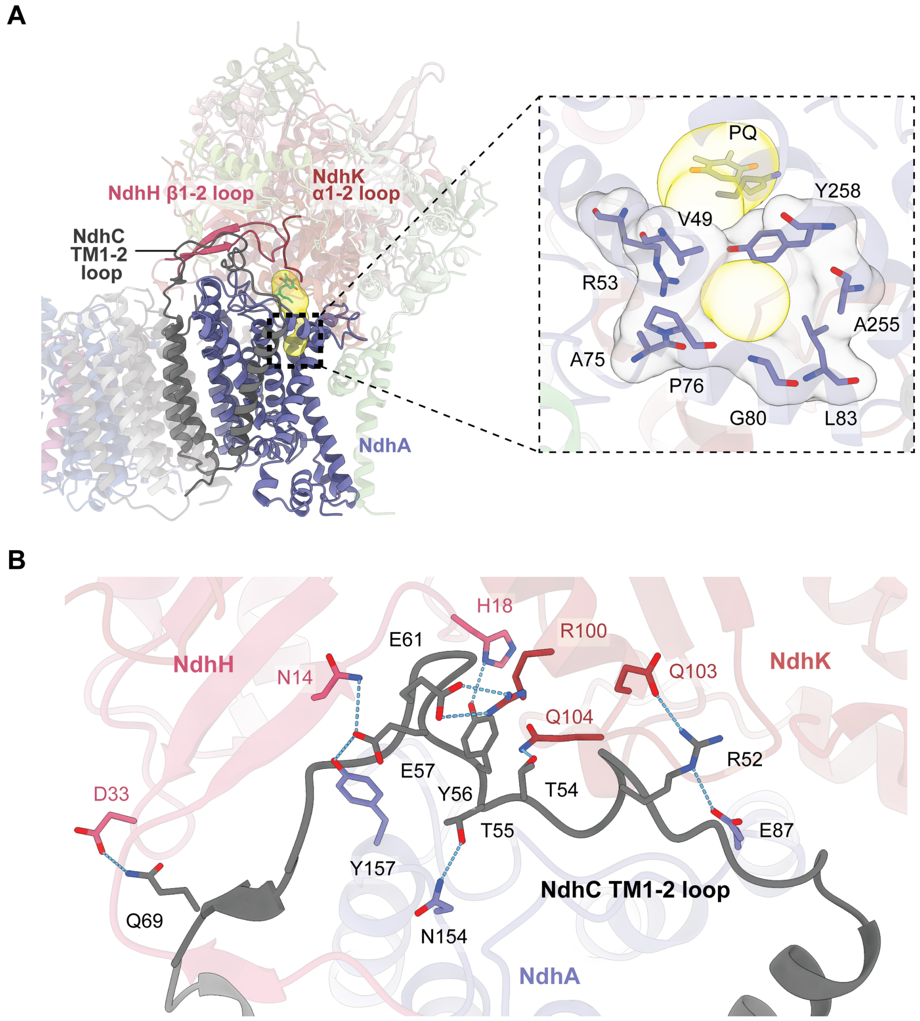

**Supplementary figure 5. Additional features of PQ chamber. A** Overview of the PQ tunnel predicted by Caver using a 1 Å radius probe (left) and zoom-in of the PQ entrance (right). The predicted tunnel is shown in yellow. **B** Hydrophilic side chain interactions with NdhH and NdhK stabilized NdhC TM1-2 loop. Hydrogen bonds and salt bridges are depicted by blue dashes.

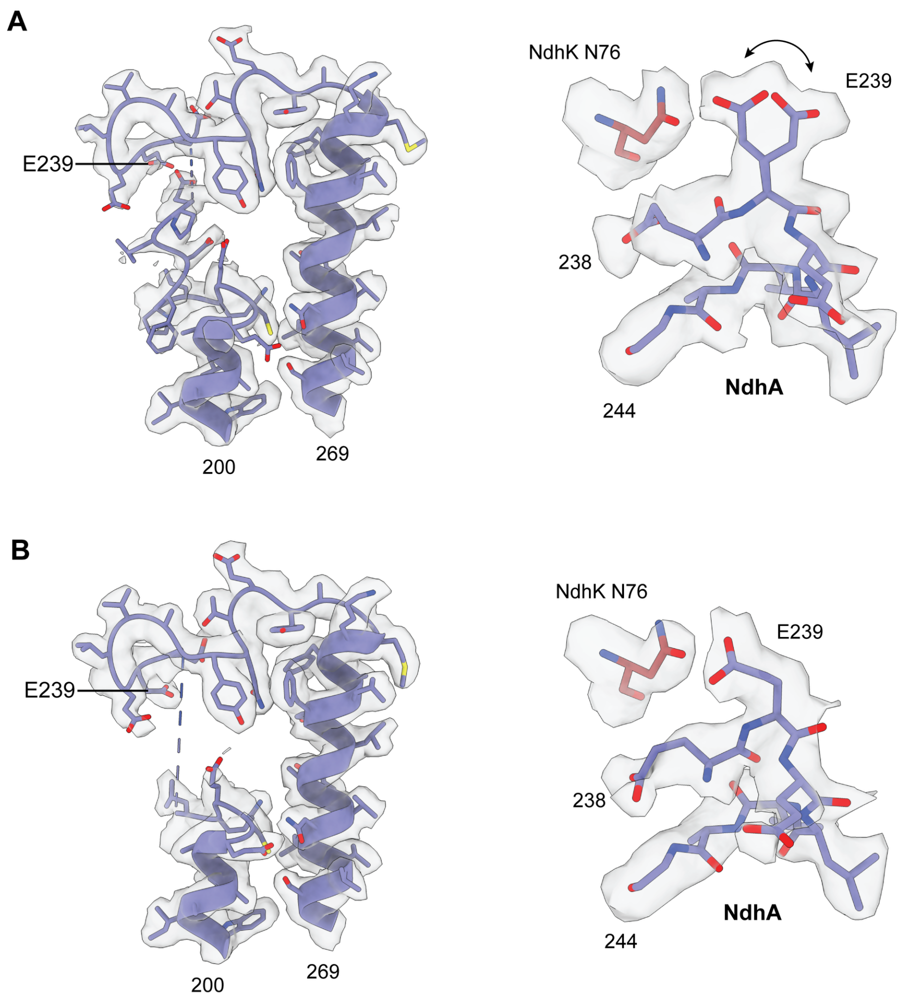

**Supplementary figure 6. Cryo-EM density of NdhA TM5-6 loop. A** Overall density fitting of TM5-6 loop (left) and local density fitting around E239 (right) in NDH-1MS' and **B** NDH-1MS'-NdhV. Notice that E239 exhibited two conformations as indicated by arrow. Densities are displayed at 4 σ. Dashes indicated unresolved regions.

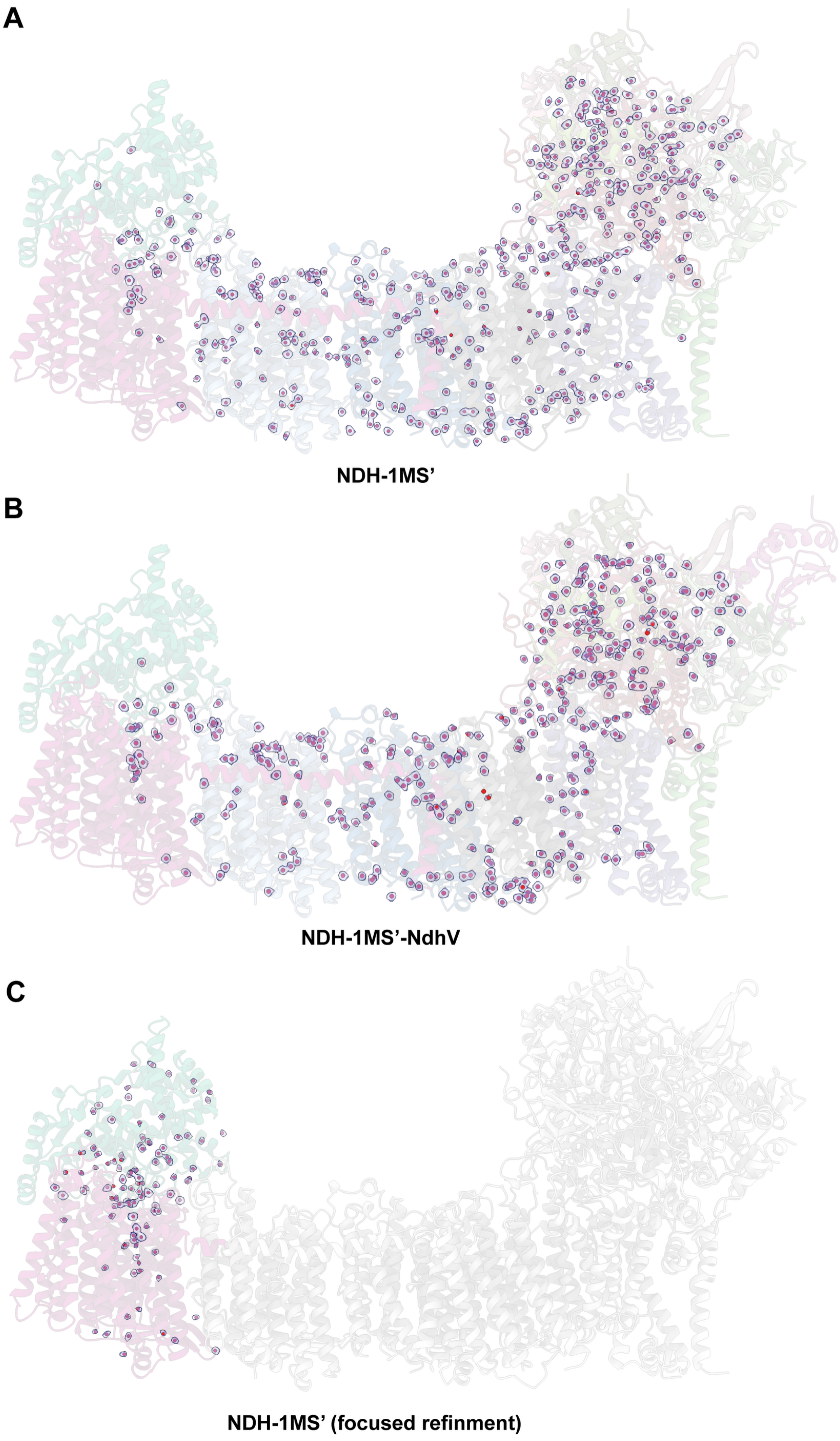

**Supplementary figure 7. Overview of structural water.** Water modeled in the consensus volume of **A** NDH-1MS’ and **B** NDH-1MS’-NdhV. Water densities are shown at 3 σ. **C** Water modeled in the focus refinement of NDH-1MS’ NdhF4-CupB module. Densities are shown at 5 σ.

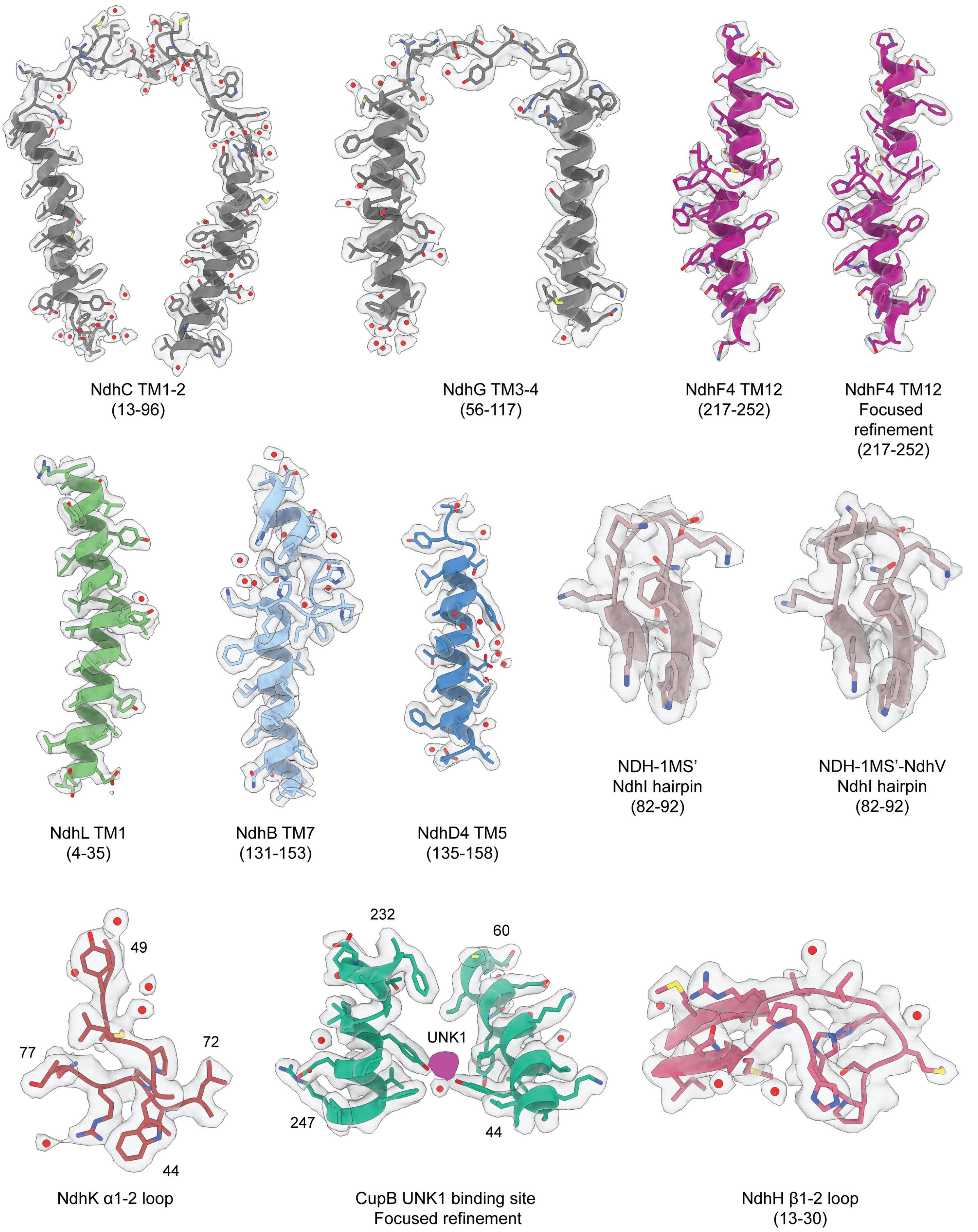

**Supplementary figure 8. Cryo-EM density example.** Representative model-density fitting of key regions in NDH-1MS'. Densities are displayed at 5 σ. For NdhI hairpin loop, densities are displayed at 2.5 σ.

### References

1. Nowaczyk, M. M. *et al.* Psb27, a cyanobacterial lipoprotein, is involved in the repair cycle of photosystem II. *Plant Cell* **18**, 3121 true 3121 3121 3131 true 3131 3131 3121–31 (2006).
2. Strohalm, M., Hassman, M., Kosata, B. & Kodícek, M. mMass data miner: an open source alternative for mass spectrometric data analysis. *Rapid Commun Mass Spectrom* **22**, 905 true 905 905 908 true 908 908 905–8 (2008).
3. Kraus, A. *et al.* Arginine-Rich Small Proteins with a Domain of Unknown Function, DUF1127, Play a Role in Phosphate and Carbon Metabolism of Agrobacterium tumefaciens. *J Bacteriol* **202**, (2020).
4. Cormann, K. U., Möller, M. & Nowaczyk, M. M. Critical Assessment of Protein Cross-Linking and Molecular Docking: An Updated Model for the Interaction Between Photosystem II and Psb27. *Front Plant Sci* **7**, 157 true 157 157 157 (2016).
5. Nakamura, Y. *et al.* Complete genome structure of the thermophilic cyanobacterium Thermosynechococcus elongatus BP-1. *DNA Res* **9**, 123 true 123 123 130 true 130 130 123–30 (2002).
6. Yates, J. R., Eng, J. K., McCormack, A. L. & Schieltz, D. Method to correlate tandem mass spectra of modified peptides to amino acid sequences in the protein database. *Anal Chem* **67**, 1426 true 1426 1426 1436 true 1436 1436 1426–36 (1995).
7. Schägger, H. & Jagow, G. Blue native electrophoresis for isolation of membrane protein complexes in enzymatically active form. *Anal Biochem* **199**, 223 true 223 223 231 true 231 231 223–31 (1991).
8. Schägger, H. & Jagow, G. Tricine-sodium dodecyl sulfate-polyacrylamide gel electrophoresis for the separation of proteins in the range from 1 to 100 kDa. *Anal Biochem* **166**, 368 true 368 368 379 true 379 379 368–79 (1987).
